# SARS-CoV-2 nucleocapsid protein engages with viral RNA and ERGIC lipids to drive viral core assembly

**DOI:** 10.64898/2026.07.23.740351

**Authors:** Shovon Swarnakar, Jhili Mishra, Virgile Rat, Peggy Merida, Louis Bunel, Damien Avinens, Jérôme Feuillard, Sébastien Lyonnais, Eric Martinez, Jitendriya Swain, Jérémy Dufourt, Mickael Blaise, Cyril Favard, Jaydeep K Basu, Delphine Muriaux

## Abstract

Severe acute respiratory syndrome coronavirus 2 assembles at the ER–Golgi intermediate compartment (ERGIC), yet the molecular basis of nucleocapsid (N) protein interactions with host membranes remains unclear. Using in vitro reconstituted lipid membranes and viral RNA– N complexes, we show that full-length N binds phosphatidylinositol (PI)- and phosphatidylserine (PS)-containing membranes and induces lipid clustering, an effect amplified by viral RNA and ERGIC-like membrane lipid composition. The isolated N-terminal domain lacks this activity, whereas the C-terminal domain retains membrane-associated multimerization. Although, lipid bilayers promote co-clustering of N and PI lipids, facilitating ribonucleoprotein (RNP) assembly, even on simple membranes. Importantly, ERGIC- mimicking membranes enhances this co-clustering further stabilizing RNPs of dimensions matching the viral core. In cells, viral RNA enhances N clustering without altering particle production. These findings reveal cooperative interactions between N, viral RNA, and ERGIC lipids as key drivers of lipid-dependent viral core formation, providing a mechanistic framework for the early steps of viral assembly.

## INTRODUCTION

Severe acute respiratory syndrome coronavirus 2 (SARS-CoV-2) is an enveloped, single-stranded positive-sense RNA virus responsible for the pandemic COVID-19 ^1,2^. A detailed understanding of the molecular mechanisms governing SARS-CoV-2 viral particle assembly remains incomplete{Citation}. Viral assembly involves the coordinated action of the four main structural proteins: the transmembrane spike (S), membrane (M) and envelope (E) proteins, located at the ER-Golgi intermediate compartment (ERGIC), and the nucleocapsid (N) protein, which packages and interacts with the viral RNA genome in the cytosol ^3–5^. Together, these proteins are sufficient to drive assembly of virus-like particles (VLPs) ^6–9^.

The Membrane M protein is the most abundant structural protein in the viral envelope and has a predominant role in virus assembly, acting as an organizing scaffold during virus formation. M is described as the interacting protein with the N protein ^10^. The E protein, an ion channel ^11^, also interacts with M and drives budding of new virus particles. The S protein incorporate the spikes into the viral envelope. The M/E proteins help the virus bud into the membranes of the host cell’s ER–Golgi intermediate compartment (ERGIC). The nucleocapsid (N) protein of SARS-CoV-2 plays a central and multifaceted role in viral assembly, linking the viral genome to the formation of mature virus particles. The N-terminal domain of N protein is made of an NTD known to bind RNA^12,13^, and a SR rich region known to be involved in N protein self- assembly^14^, its Cter-CTD domain has also been described to be involved in N protein self-assembly and RNA binding^15^ and, more recently, in lipid binding ^16^.

The primary function of the cytosolic N protein is to bind the viral positive-sense RNA genome, of about 30kB, through an N-terminal RNA-binding domain^17,18^ and a C-terminal dimerization domain^12,13^. Multiple N proteins coat the viral RNA genome, forming a flexible ribonucleoprotein (RNP) complex, therefore compacting and protecting the genome for packaging inside the nascent viral particle. N proteins self-assemble (dimerize and oligomerize), multimerize on the viral RNA genome to be organized into higher-order structures, named the viral core or RNP (RiboNucleoComplex)^19^. The RNP buds from the ERGIC membrane, enabling acquisition of a host-derived lipid envelope enriched in M, E and S proteins. The mechanism of how the assembly of the RNP occurs on M/E and S and at the ERGIC membrane remains poorly described. A direct or indirect interaction of N protein with the membrane (M) protein has been proposed ^20^ and that the M–N interaction recruits the RNP complex to the budding site, where virion assembly takes place. Unlike rigid viral capsids, the N–RNA complex is supposed to be structurally flexible, which fits the pleomorphic (variable) shape of coronaviruses and allows efficient packaging of the large (∼30 kb) genome. In summary, the SARS-CoV-2 N protein packages and protects the viral RNA, self-assembles into ribonucleoprotein (RNP) complexes, and interacts with the M protein to ensure that the genome is incorporated into budding virions. It is proposed that M acts as a critical bridge between the viral genome and the structural assembly machinery. In the case of SARS-CoV-2 Virus-like particle (VLP) formation, it was reported that the minimal structural viral proteins to build up a VLP are M, E and N or S; in the absence of one of these proteins, no VLP are produced ^6–8,21^.

Thus, a direct protein-protein interaction between M and N at the ERGIC membranes might not be the only favorable mechanism to ensure packaging of the viral genome inside the virion. Interestingly, N has been recently shown to interact with anionic lipids with an implication of the C-terminal domain of N ^16^, but no description has been provided for a possible role of N assembly on ERGIC membrane and in modulating/assisting RNP complex assembly.

Here, we sought to clarify how the N protein—and the RNP complex—interacts with lipid membranes. Using in-vitro approaches with simplified ^22,23^ and ERGIC-mimicking lipid membranes ^24^, we examined the behavior of N and pre-assembled RNPs on single lipid bilayer mimicking membranes, proposing the hypothesis that an intermediate N-lipid, or RNP-lipid, interaction exists early in assembly and can promote RNP assembly on ERGIC membrane. Finally, we evaluated the impact of the viral RNA on intracellular RNP generation, and on the incorporation of N in Virus-like Particles (VLP) as compare to natural extracellular vesicles (EV) in pulmonary cells.

Our results show that the N protein can bind large unilamellar vesicles (LUVs) composed of PC:PS and PI, an ERGIC lipid, albeit with low affinity and without specific preference for PIP derivatives. Notably, N can induce clustering of PI (and PS) lipids within biomimetic membranes, an effect amplified by the presence of viral RNA and ERGIC-like lipid compositions of the Supported Lipid Bilayer. This lipid clustering property of N was dependent on the C-terminal domain of N. In the presence of viral RNA, RNP complexes can form, and the addition of a lipid bilayer further promotes the co-assembly of lipids and N protein, facilitating RNP formation in vitro. On ERGIC-mimicking membranes, RNP assembly is enhanced, promoting the formation of an RNP complex with dimensions consistent with the SARS-CoV-2 viral core. In pulmonary cells, we similarly observe that intracellular RNP clustering is improved in the presence of viral RNA, generating structures comparable in size to the viral core, and that can meet ERGIC membrane.

Altogether, these findings provide new insights into how the SARS-CoV-2 core assembles on ERGIC lipid membranes to drive viral assembly, and propose a model for how the viral RNA and the PI lipids can facilitate SARS-CoV-2 core assembly at ERGIC membranes.

## RESULTS

Although SARS-CoV-2 nucleocapsid (N) protein has been shown to interact with anionic lipids in vitro ^25^, it remains unclear whether the viral ribonucleoprotein (RNP) complex formed by N and the viral RNA exhibits specificity for particular membrane lipids, or whether distinct lipids simply serve as anchors for N to initiate the assembly process. In this study, we first investigate the assembly of the N protein, both in the absence and presence of viral RNA, on *in vitro* model membranes and within cells producing N-containing virus-like particles.

### Purified recombinant SARS-CoV-2 N proteins and RNP in buffer solution

We first study N and the viral ribonucleocomplex (RNP), made of in vitro-synthesised and purified N with a pseudo-viral RNA, in buffer solution (Figure 1). SARS-CoV-2 (Wuhan strain) N proteins, WT or truncated N-ter or C-ter N proteins, were generated and purified in vitro with or without His-Tag (See Material and Methods, Figure 1 A-B and Figure Supplemental 1 A-B for further details of protein purification). To mimic the viral RNA, we generated, using an *in vitro* transcription assay, an RNA made of the SARS-CoV-2 viral 5’UTR and 3’UTR with a synthetic sequence in between to get a 2500nt-long viral RNA (see Material and Methods), named vRNA (Figure 1C-D).

**Figure 1:**
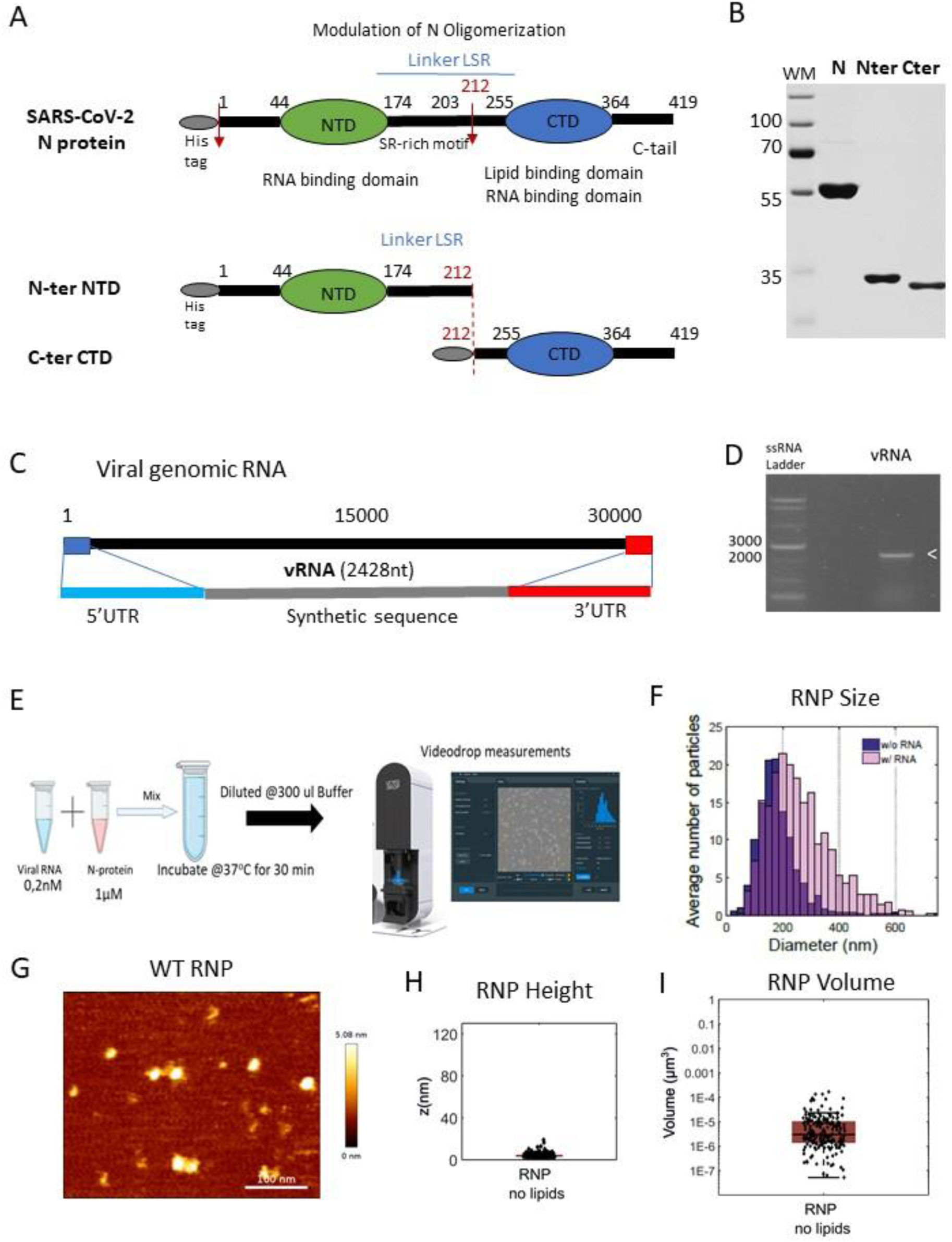
SARS-CoV-2 N proteins, viral RNA and viral RNP in buffer solution. **A**) Scheme of the WT N and N-ter, C-ter truncated N protein of SARS-CoV-2 used in this study. SARS- CoV-2 N protein domains with the His-tag domain and the cleavage site. Scheme of the Nter- NTD (1-212) and Cter-CTD (212-419) domains of purified truncated N proteins. **B**) Coomassie stained acrylamide gels showing purified N, N-ter and C-ter proteins used in this study. WM is the protein ladder for sizes. **C)** Scheme of the Synthetic pseudo-viral RNA (2428 nt long) used in this study. **D**) Agarose gel showing the in vitro transcribed purified viral RNA. **E)** Scheme of the Videodrop measurements. **F**) Distribution size measurement of WT N (in nm) in buffer solution in the absence (violet color) or in the presence of viral RNA (pink color), as indicated. N=4 independent experiments with 2 different lots of purified WT N proteins. **G**) Image of RNP deposit on a coverslip using AFM. **H-I**) Measurements of RNP height (z in nm) and volume (in µm3) in the absence of lipids, using AFM.

N proteins (1 µM) were incubated in physiological buffer (150 mM NaCl, 10 mM Tris– HCl, pH 7.4) with or without vRNA (2nM) for 30 min at 37°C, and the mixture was further analysed for RNP formation either in solution, using a Videodrop, allowing for quantification of particle diameter and concentration (Figure 1 E-F) or deposited on a glass surface using Atomic Force Microscopy (AFM). Our results show that incubation of the N protein with the viral RNA in buffer solution triggers the formation of RNPs with a diameter ranging between 100 and 500nm (242+/-153 nm, median+/-IQR), while in the absence of RNA, N self-assembles in a smaller complex lower than 200nm diameter (168+/-79 nm) (Figure 1F). We then deposit these RNPs on a glass coverslip and use AFM to image the RNPs (Figure 1G-H, Supplemental Figure 2A). We observe an RNP height of 6 nm on average, with a median volume of 3.10e^−5^ µm^3^ in the absence of lipids (“no lipid”). These RNPs observed by AFM are much smaller units (median height 6 nm and median diameter size 20 nm) than the RNP size distribution observed in solution (up to 400nm), indicating that the higher order RNPs seem quite unstable in buffer solution and are made of small stable 20nm length units when deposited on a surface.

### Screening for Lipid interactions with N proteins on LUV

We used liposome co-sedimentation assays and large unilamellar vesicles (LUVs) (Figure 2) with different lipid compositions to identify the specific lipids that are critical for N protein interaction under physiological salt concentration (10mM Tris, 150 mM NaCl, pH=7.4). LUVs with minimal lipid compositions found in ERGIC membrane, like PC, PS and PI, were prepared and incubated with N protein in the presence or in the absence of viral RNA (Figure 2).

**Figure 2:**
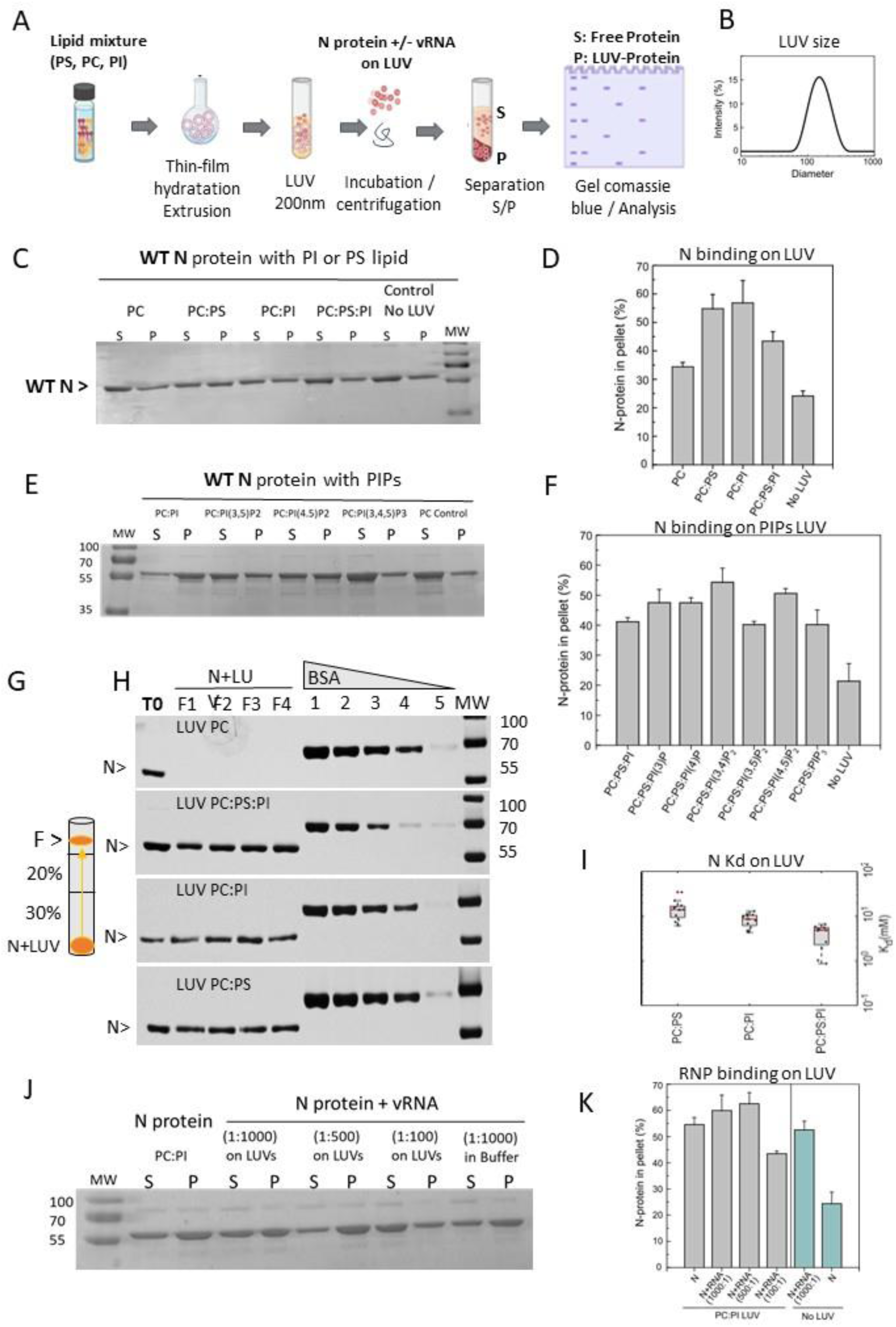
SARS-CoV-2 N proteins or viral RNP interaction with LUV of different lipid compositions. **A**) Scheme of the LUV co-sedimentation assay and experimental procedure; **B**) Size of the LUV (200nm in diameter) used in this study, as measured by Dynamic Light Scattering (DLS); **C**) Western blot of WT N protein interaction with LUV of simple composition mixing PC:PS:PI lipid at different ratio, as indicated; **D**) quantification of N protein-LUV binding by mean of the Percentage of N found in the Pellet after centrifugation over the total N fractions (P+S); **E**) Western blot of N protein interaction with different phospholipids (PIPs) on LUV; **F**) Quantification of the % of N binding to the LUVs : %P=P/(P+S)x100; **G**) Scheme of the N-LUV flotation assay principle; A mix of N+LUV is loaded at the bottom of the tube. F is the fraction containing N binding to the floating LUV; **H**) Example of N LUV flotation assays on LUV PC and PC:PS:PI (80:10:10). To calculate the apparent Kd, a BSA gamme has been loaded on each SDS-PAGE gel. **I**) Measurement of the apparent affinity constant of the N-protein depending on the LUV lipid compositions by float- up experiment. For each lipid composition, three independent experiments (N=3) with four Kd measurements each (n=4) were performed. **J**) Western Blot of [N+vRNA] on LUV at different Protein:RNA ratio, as indicating; **K**) Quantification of vRNP binding on LUV and found in the Pellet. S = N protein in the Supernatant and P = N protein in the Pellet after centrifugation. All the Western Blot are revealed with a SARS-CoV-2 anti-N protein. N=3 to 4 independent experiments. P: pellet fraction; S: supernatant fraction after centrifugation.

After incubation of N with 200nm diameter LUV at 37°C for 30 min, samples were centrifugated to separate the supernatant (S) fraction containing the unbound soluble N from the pellet (P) fraction containing the LUV-bound N (Figure 2A-B). N protein incubation without LUVs served as a negative control to estimate the fraction containing N aggregates in the pellet. Both fractions were then migrated on SDS gel and stained with Coomassie blue for N visualisation and quantification. It was observed that 20% of the N protein was without LUV aggregates in the pellet and represents the fraction of insoluble N. It was found that 30% of N protein pellet with PC only, but 55 to 60% binding with PC: PS (8:2 mol: mol) or PC: PI (8:2 mol: mol), or PC:PS:PI (8:1:1 mol:mol:mol) was observed, indicating a preference of N for anionic charged PI-containing LUV (Figure 2C-D). Our results showed that N binds with PI-containing lipid mixtures, a predominant ERGIC lipid. Possibly His-Tag effect on the N-lipid binding efficiency and specificity was discarded (Supplemental Figure 3A-B).

As it was reported that N is binding to other PIP derivatives on Lipid Strips ^25^, we explore the binding of N on PIP-LUV using PI derivatives like PI(3)P, PI(4)P, PI(3,4)P2, PI(3,5)P2, PI(4,5)P2, PI(3,4,5)P3: we found that N proteins mostly bound significantly with negatively charged PI lipid and its PI derivatives without any significant differences (Figure 2E-F), suggesting that N could bind any cellular membranes as long as it is anionic and accessible but with a preference for PI/PS, as main phospholipids of the ERGIC membranes, the site of the virus assembly.

We thus determine the apparent K_d_ for N binding to PI, PS or PS:PI:PC containing LUV. N- protein affinity for membranes was measured by performing float-up experiments in which N- protein was mixed with large unilamellar vesicles (LUVs), and the N-protein–bound fraction was separated from the free fraction using a density gradient (Figure 2 G). LUVs were mixed with N-protein to reach a final lipid concentration of 5 mM and N-protein concentration of 3 µM. The protein–LUV solution was incubated at room temperature for 30 minutes to allow equilibration. Following incubation, the LUV–protein solution was mixed with OptiPrep density gradient to proceed for ultracentrifugation. After ultracentrifugation, the floated LUVs were collected in the presence of 0.5 mol% C18 DiI lipids (Figure 2G). After collection, the N- protein content in the floated fraction and the initial protein concentration before flotation were quantified by PAGE gel electrophoresis (Figure 2H). Protein concentration was measured on the gel using a BSA titration standard curve, providing the bound protein concentration [P]_bound_ and the total protein concentration [P]_total_. The dissociation constant Kd of the N-protein (Figure 2I) was found to be 14.95 ±7.65, 8.45 ±2.69 and 4.14 ±1.97 µM on PC:PS, PC:PI and PC:PS :PI LUV, respectively, showing a significantly higher affinity of N for PI:PS containing LUV.

Further, we tried to explore N binding to the PI containing LUV in the presence of N pre-mixed with the vRNA, but it was difficult to observe any significant enhancement of the RNP binding since the RNP complex already sedimented in the absence of LUV (Figure 2 J-K). In order to circumvent this issue, we decided to study N and RNP assembly on SLB membranes.

### RNA enhances Phospholipid clustering by the SARS-CoV-2 N proteins on simple and ERGIC-like membranes

In order to study the assembly of N, in the absence or in the presence of the vRNA, on lipid membranes, we performed phospholipid clustering assays on supported lipid bilayers (SLB) of different lipid composition, either simple (PC:PS:PI, 80:10:10, mol:mol:mol) or more complex supported lipid bilayers (SLB), mimicking the ERGIC (PC:PE:PS:PI:Chol (50:20:10:10:10), ie. the SARS-CoV-2 assembly membrane (Figure 3A). To further analyse the interplay between N, RNP and the specific phospholipids PI and PS recognised by N, we monitored the ability of N or RNP to cluster fluorescently labelled PI or PS lipids on the different SLB biomimetic membranes (Figure 3 and Supplemental Figures 4 and 5).

**Figure 3:**
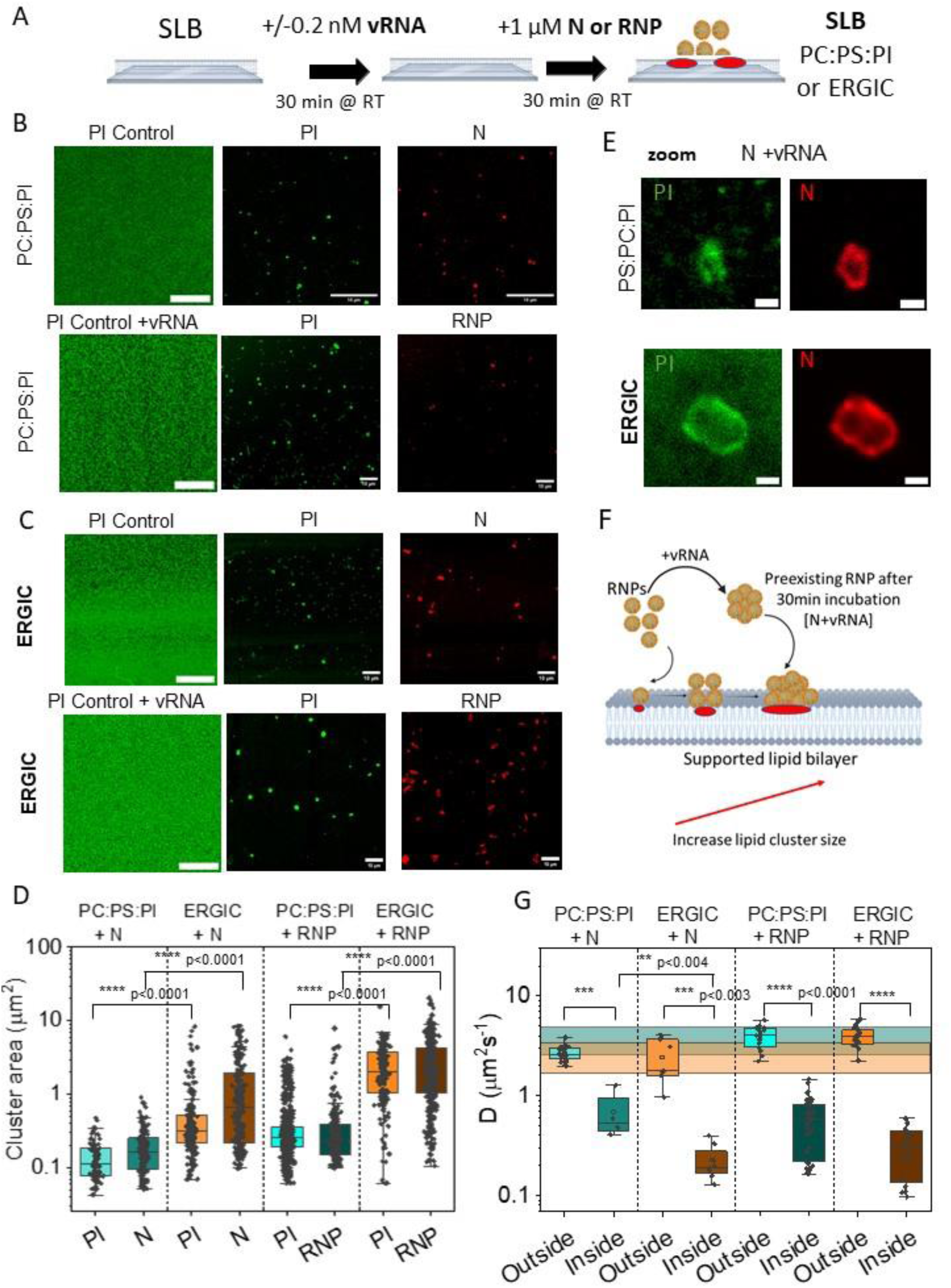
SARS-CoV-2 N Protein and RNPs Induce Phosphatidylinositol (PI) Clustering and Trapping on simple and ERGIC Biomimetic Membranes. **A)** Scheme of the experimental design. **B)** Imaging SARS-CoV-2 N protein-PI clusters on lipidic biomimetic membranes of simple and ERGIC-like composition using confocal microscopy. Confocal imaging showing WT-N selectively generates fluorescent TopFluor-PI lipid clusters (in green) on SLBs of basic lipid composition made of PC:PS:PI (80:10:10) and on **C)** ERGIC-like composition made of [PC:PE:PS:PI:Chol (50:20:10:10:10)], N and PI clusters induced by N are visible on the SLBs. SARS-CoV-2 N+viral RNA (vRNA) complex enhances PI lipid cluster formation on SLBs of basic lipid composition and ERGIC-like composition are shown. 1 to 5% of the N protein was labelled with N-Alexa647 (in red). Scale bar is 10µm. **D**) Graph showing the quantification of PI and N cluster areas (in µm^2^) in the presence of N protein or RNP (N+vRNA) on simple PC:PS:PI SLBs or on ERGIC-like membranes (also reported in Table 1). **E**) Imaging TopFluor-PI lipid clustering (in green) induced by N proteins on PC:PS:PI or ERGIC membranes in the presence of viral RNA (vRNA), using confocal microscopy. Scale bar is 20µm. **F**) Model of RNP formation and PI lipid clustering on SLB. **G**) Box plots showing the diffusion coefficients (*D*) of TopFluor PI lipid inside and outside the PI clusters induced by the N protein or RNP (N+vRNA) on simple PC:PS:PI SLBs or on ERGIC-like membranes using FCS. The cyan and brown patch showing the control diffusion coefficient for PC :PS :PI and ERGIC-like membrane respectively. An average of 100 to 900 clusters were analyzed for each condition (from N=3 independent experiments). For FCS data, an average of 5 to 45 *D* values is presented. Data are presented as box-and-whisker plots, showing the median, 1st and 3rd quartiles. Differences were evaluated using the Mann-Whitney test (p values are indicated).

**Table 1:**
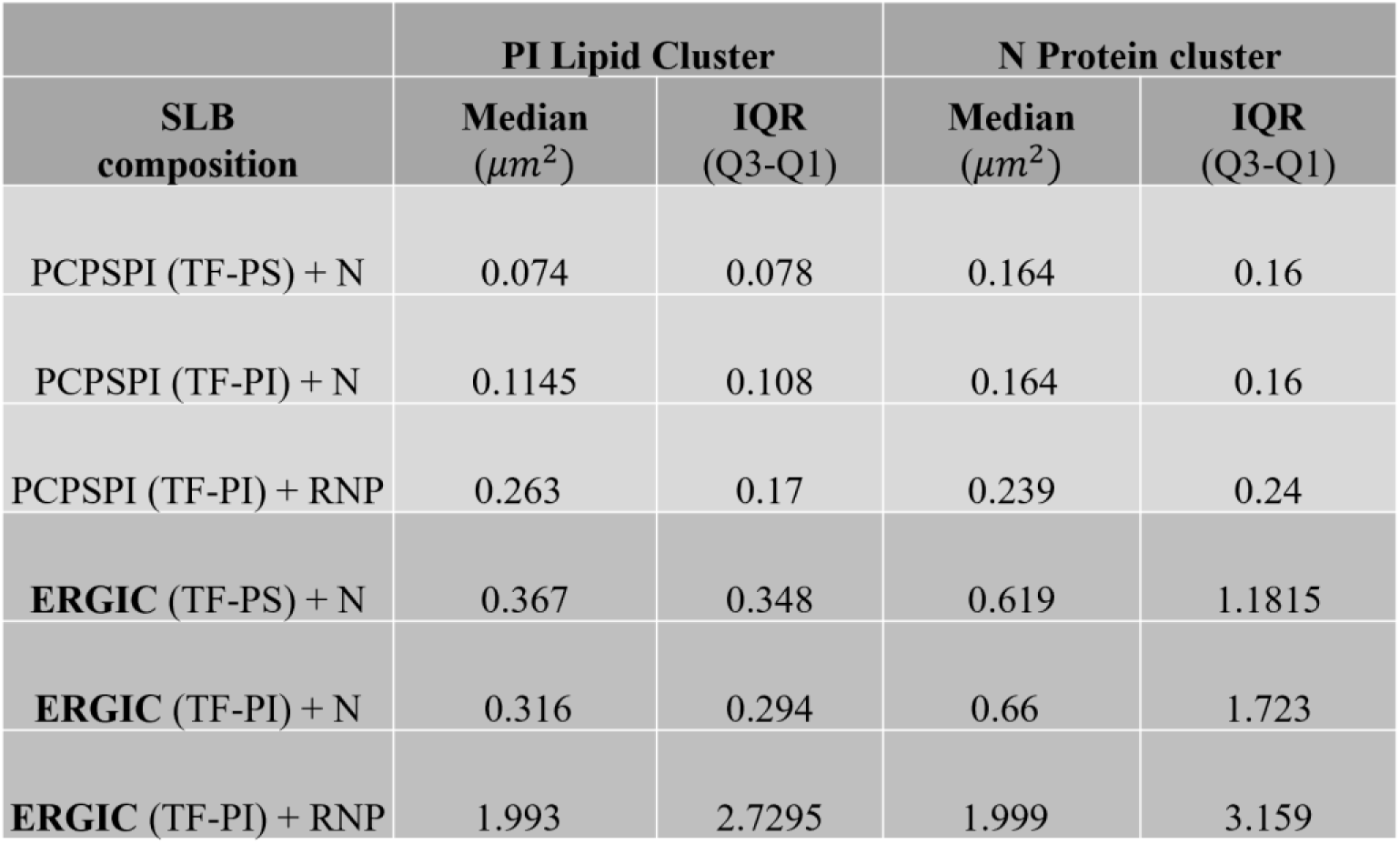
Lipid and N protein cluster sizes upon N multimerisation on SLB using confocal microscopy for fluorescence measurements – Labelled PI or PS lipids are estimated at 1 to 5% of labelled N. Purified N was cleaved of the His tag. Data measurements are Median ± IQR (Q3 upper quartile 75%-Q1 lower quartile 25%).

| SLB composition | PI Lipid Cluster |  | N Protein cluster |  |
| --- | --- | --- | --- | --- |
| | Median ($\mu\text{m}^2$) | IQR (Q3-Q1) | Median ($\mu\text{m}^2$) | IQR (Q3-Q1) |
| PCPSPI (TF-PS) + N | 0.074 | 0.078 | 0.164 | 0.16 |
| PCPSPI (TF-PI) + N | 0.1145 | 0.108 | 0.164 | 0.16 |
| PCPSPI (TF-PI) + RNP | 0.263 | 0.17 | 0.239 | 0.24 |
| **ERGIC** (TF-PS) + N | 0.367 | 0.348 | 0.619 | 1.1815 |
| **ERGIC** (TF-PI) + N | 0.316 | 0.294 | 0.66 | 1.723 |
| **ERGIC** (TF-PI) + RNP | 1.993 | 2.7295 | 1.999 | 3.159 |

We did not detect any lipid cluster on SLB in the buffer or in the presence of the vRNA alone (the median lipid cluster size was found to be 0.07 µm², which corresponds to the diffraction limit detection in the confocal fluorescence microscope for a 488 nm excitation (∼260 nm)). In the presence of N or N + vRNA (RNP), after 30 min of incubation either at 37°C or at RT, we observed PI cluster formation on PI-TF containing basic SLB (Figure 3B) or ERGIC SLB (Figure 3C) . PI clustering similarly occurs in the presence of another probe like PI-TMR with N (supplemental Figure 4). Area of the PI-TF lipid cluster generated by N after 30 min of incubation on basic and ERGIC SLB was measured (Figure 3D) and is recapitulated in Table 1 (Median ± IQR). In some experiments, a fraction (1 to 5%) of the protein N was labelled with Alexa-647 (see Methods), and N clustering was also evaluated on the different SLBs and the area cluster sizes measured (see Table 1). We measured small lipid clusters induced by N binding for PS or PI (0.07-0.11µm^2^) on PC:PS:PI membranes, whereas, in the presence of the viral RNA, the lipid cluster size increased by 2-3-fold, suggesting an improved clustering in the presence of viral RNA. Importantly, fluorescently labelled N cluster sizes closely match the lipid cluster sizes, reinforcing the idea that the lipid clusters are induced by the presence of N protein with or without RNA. On the ERGIC biomimetic membrane, the PI (or PS) lipid cluster sizes were found 3-4-fold bigger (0.32-0.37 µm2) in the presence of N alone and increased again by 5-6-fold upon addition of the viral RNA (1.99 ±2.72 µm²). We check that the PI clustering was not dependent on the fluorescent label borne by the lipid probe. Indeed, the clustering of TMR-PI was also enhanced by the RNP as compared to N alone on PC:PS:PI membranes (Supplemental Figure 4). Due to the partial labelling of N, we could prove that the PI or PS clustering on SLB indeed contained the N protein (Figure 3B, Supplemental Figure 4C and 5B).

Our results indicate that the clustering of fluorescent labelled PI or PS lipids is induced by the viral N protein on the lipidic membrane and is furthermore enhanced by the viral RNA. The lipid clustering beneath assembled RNP on the ERGIC membrane is the most efficient.

Furthermore, to get insight into this lipid clustering, we characterised the mobility of these fluorescent PI lipid analogues (TF-PI) within and outside the N-induced PI clusters using, a widely used technique Fluorescence Correlation Spectroscopy (FCS)^26–28^ to study single molecule mobility on both PC:PS:PI and ERGIC-mimetic membranes. The distribution of lipid diffusion coefficients (D) (Figure 3G) indicates that lipid dynamics within the lipid clusters are significantly constrained compared to the surrounding membrane, where mobility remains comparable to the control values. While the diffusion coefficient of TF-PI in SLB was relatively similar (D = 2.90 ± 0.44 μm^2^s^−1^ for PC:PS:PI and D = 1.79 ± 0.46 μm^2^s^−1^ for ERGIC). Before the injection of N, the impact of the N protein differed between the two systems. Specifically, the reduction in lateral mobility was more pronounced in the ERGIC-mimetic membrane, where diffusion slowed approximately 9-10-fold (D = 0.19 ± 0.11 μm^2^s^−1^), compared to a 4-5-fold reduction in the PC:PS:PI membrane (D = 0.54 ± 0.49 μm^2^s^−1^). Similarly, lipid trapping was observed in the presence of the RNP complex. The difference between the D values inside and outside PI cluster dynamics was further amplified in the presence of the RNP complex, showing a 7-fold reduction in PC:PS:PI (D = 0.58 ± 0.60 μm^2^s^−1^) and a 13-fold reduction in the ERGIC membrane (D = 0.30 ± 0.31 μm^2^s^−1^) (Table 2). This suggests that the N protein or the RNP sequesters PI lipids more efficiently within the ERGIC environment.

**Table 2:** TF-PI diffusion D values outside or inside N clusters on PC:PS:PI or ERGIC SLB in the presence of N proteins or RNP. D values are represented as the Median±IQR (Q3 upper quartile 75%-Q1 lower quartile 25%).

| System | PI Lipid diffusion outside<br>($\mu\text{m}^2\text{s}^{-1}$) | | PI Lipid diffusion inside<br>($\mu\text{m}^2\text{s}^{-1}$) | |
| --- | --- | --- | --- | --- |
| | Median<br>($\mu\text{m}^2$) | IQR<br>(Q3-Q1) | Median<br>($\mu\text{m}^2$) | IQR<br>(Q3-Q1) |
| PCPSPI (TF-PI) + N | 2.56 | 0.63 | 0.54 | 0.49 |
| PCPSPI (TF-PI) + RNP | 4.03 | 1.67 | 0.58 | 0.60 |
| ERGIC (TF-PI) + N | 1.77 | 2.12 | 0.19 | 0.11 |
| ERGIC (TF-PI) + RNP | 3.89 | 1.271 | 0.30 | 0.30 |

Collectively, these results demonstrate that N-lipid interactions are significantly enhanced in the ERGIC membrane composition and that the presence of the viral RNA, i.e.. The RNP complex enlarges the surface of interaction with the membrane, clustering a higher number of lipids. Notably, RNP–lipid interactions mediated by N and the viral RNA on membranes led to the formation of higher-order area (>20 µm²), ring-like structures, in approximately 20% of the clusters (Figure 3E). These structures were smaller on PC:PS:PI membranes than on ERGIC membranes and resembled the multimerisation of N on circularised viral RNA (Figure 3E).

### The C-ter domain of the N protein is responsible for PI lipid clustering on membranes

To identify if the N-ter or the C-ter domain of the N protein was required for lipid clustering on membranes *in vitro*, we produced and purified the N-ter (1-212) and C-ter (213-419) domains of N (Figure 1A and B, Supplemental Figure 1C-D). We first check in the buffer solution if, in the presence of vRNA, RNP is forming. When incubated, the N-ter or the C-ter of N were unable to form any detectable RNP in solution as measured by the Videodrop (Figure 4A), either because there is no RNP forming or because they are smaller than 80nm in diameter, thus undetectable by the Videodrop.

**Figure 4:**
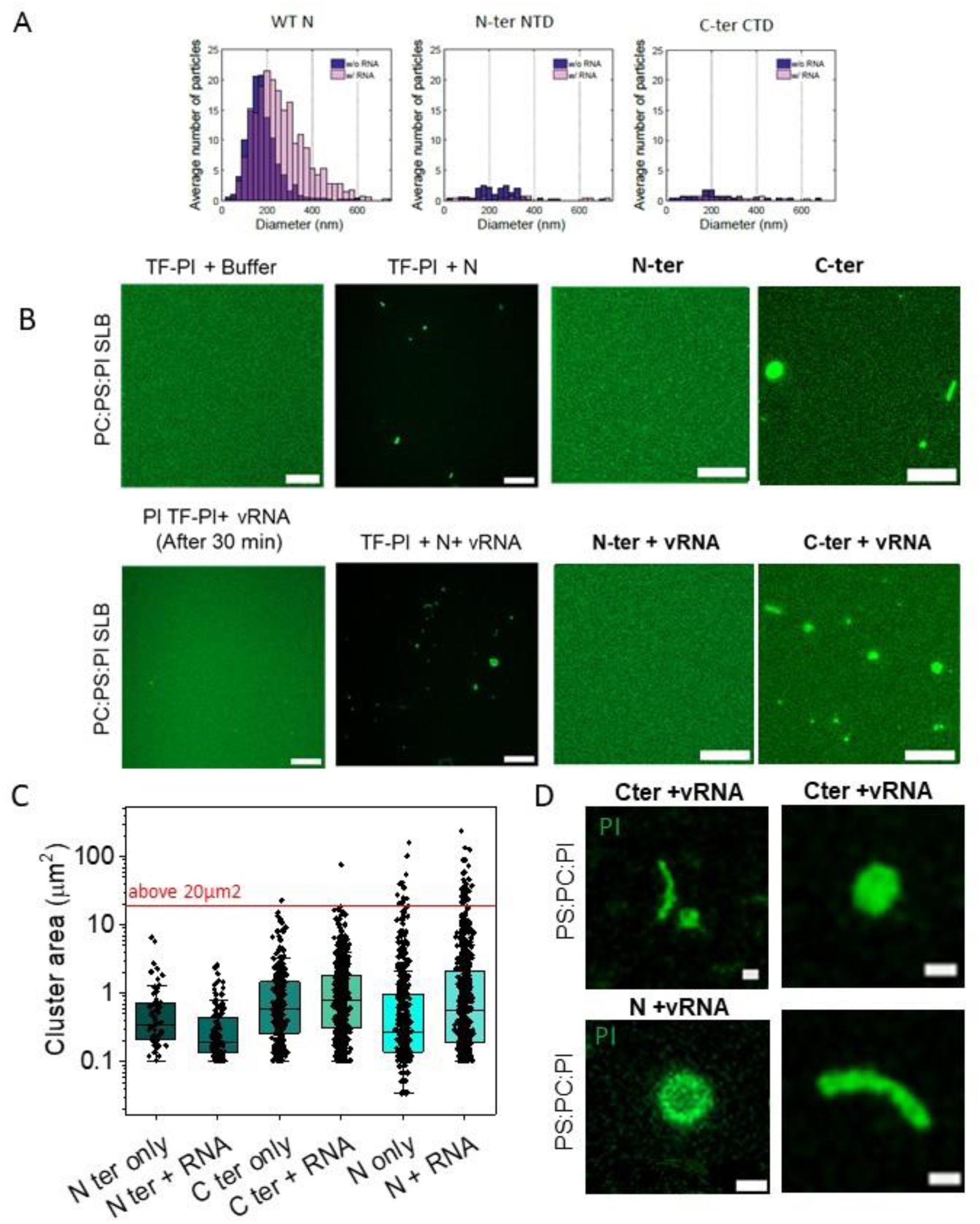
PI-TF lipid clustering on PC:PS:PI SLB comparing N-ter and C-ter proteins with WT N with and without viral RNA. **A**) N and RNP measurements using the Videodrop: WT N, N-ter and C-ter domains with or without addition of 2nM of viral RNA in buffer solution without lipids; **B**) Confocal imaging TopFluor-PI clustering (in green) induced by SARS-CoV- 2 N, N-ter or C-ter proteins on PC:PS:PI (80:10:10) biomimetic membranes, without or with viral RNA (+vRNA). All proteins are HIS-tagged proteins. **C**) Graph showing the quantification of PI cluster areas (in µm^2^) in the presence of different N, N-ter, C-ter proteins + or - vRNA, as indicated (values are also reported in Table 2). **D**) Zoom imaging of TF-PI lipid clustering (in green) induced by C-ter protein with vRNA on PC:PS:PI membranes showing the formation of filaments and filled clusters as compared to N+vRNA forming rings. Scale bar is 10µm. (all proteins are HIS-tagged proteins) **A**) Size distribution of RNP made of N+vRNA or Nter or Cter measured by Videodrop **B**) Confocal imaging TopFluor-PI clustering (in green) induced by SARS-CoV-2 N, N-ter or C-ter proteins on PC:PS:PI (80:10:10) biomimetic membranes. **C**) with the viral RNA (vRNA). **C**) Graphe showing the quantification of PI cluster areas (in µm^2^) in the presence of different N, N-ter, C-ter proteins + or - vRNA, as indicated (values are also reported in Table 2). **D**) Zoom imaging of TF-PI lipid clustering (in green) induced by C-ter protein with vRNA on PC:PS:PI membranes showing the formation of filaments and filled clusters Scale bar is 10µm.

**Figure 5.**
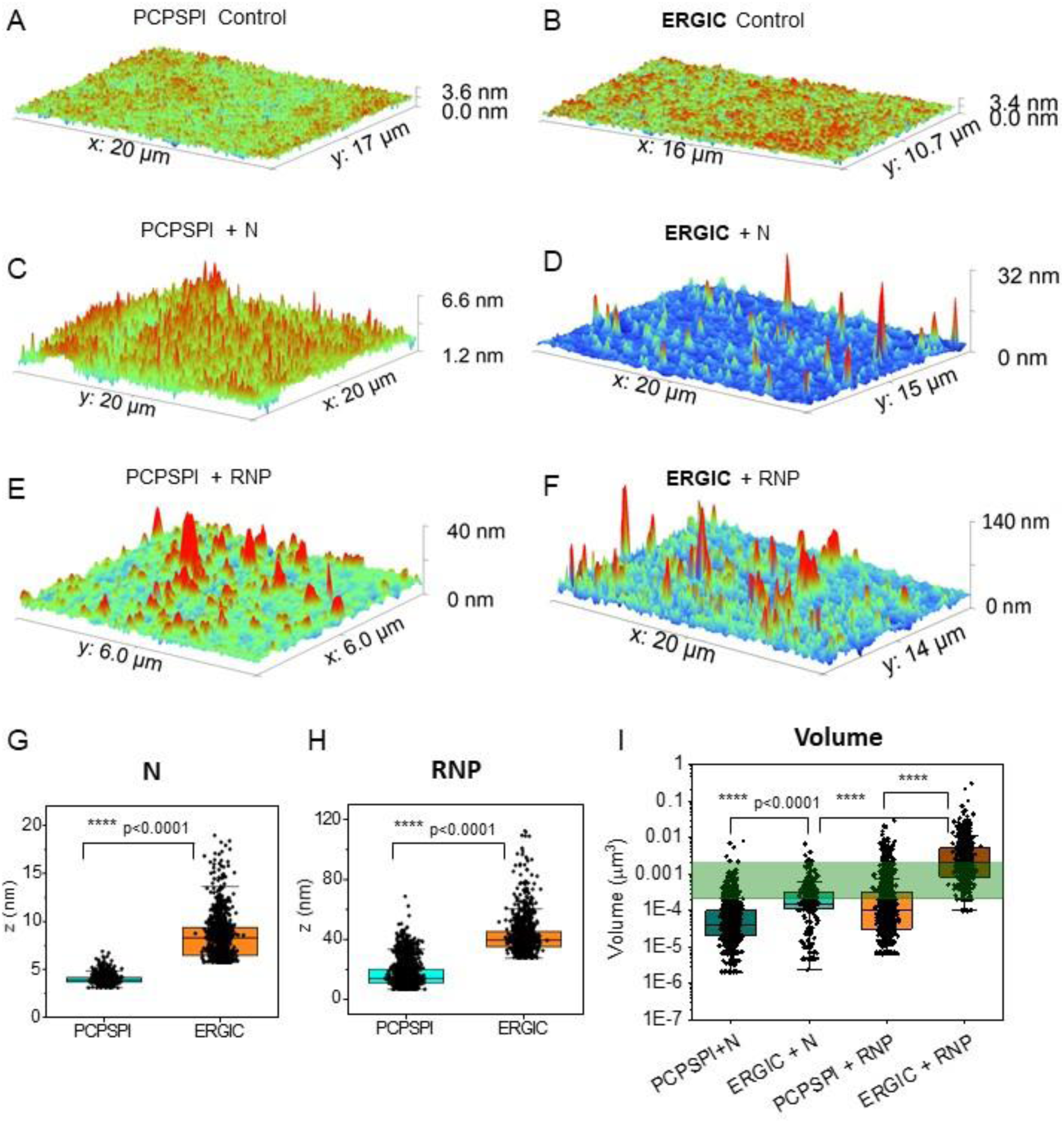
Morphology (size and volume) of SARS-CoV-2 N protein-lipid clusters on lipidic biomimetic membranes using Atomic Force Microscopy. The experiments were done in mirror of the Figure 3, SLB in the presence or absence of viral RNA, on basic (PC:PS:PI) and ERGIC mimicking bio-membranes on which 1µM N protein is deposit for 30min. **A-C)** Showing a 3D representation of AFM images on A) control PCPSPI membrane **B)** 30 min after the N protein and **C)** RNP clusters. Similarly, **D-F)** showing for **D)** control ERGIC mimicking membrane **E)** 30 min after the N protein and **F)** RNP clusters**. G)** The box plots for height (Z) comparison of N-lipid cluster between PCPSPI and ERGIC mimicking membrane and **H)** showing Z comparison of RNP-lipid cluster between PCPSPI and ERGIC mimicking membrane. **I)** The box plots for volume (V) comparison of N/RNP-lipid cluster between PC:PS:PI and ERGIC-like membrane. The 3D plots, Z and V estimations were done using Gwyddion software. The Green patch showing the typical size of a SARS-CoV-2 virion. Data are presented as box-and-whisker plots, showing the median, 1st and 3rd quartiles. An average of 960 clusters were analyzed for 4G, 1080-2900 for 4H and 360-1680 clusters for 4I (from N=2 independent experiments). Differences were evaluated using the Mann-Whitney test (p values are indicated).

We then investigate if the N-ter or the C-ter of N were able for PI-TF clustering on PC:PS:PI membranes and evaluate individual N-ter (1-212) and C-ter (213-412) domains in this assay in the absence or presence of the viral RNA (Figure 4B). We observe no lipid clustering in the presence of N-ter with or without RNA (Figure 4B), suggesting that the N-ter alone is not sufficient to induce PI lipid clustering on a membrane. The most striking was the remaining capacity of the C-ter to cluster lipid on an SLB (Figure 4B) as RNPs made by C-ter were not detectable in solution (Figure 4A). The efficacy of the C-ter domain, known to contain N multimerisation domains ^25^, at inducing PI lipid clustering was superior to N but not in generating high-order RNP (above 20µm2) (Figure 4C). We report here the density of clusters (per frame of imaging). While N-ter with or without viral RNA shows no cluster formation (median = 4.41e-4 ± 1.65e-4 clusters per µm^2^ ), we measured that the density of PI cluster lipid induced by Cter alone is almost 2-fold higher (7.72e-4 ± 2.48e-4 clusters per µm^2^ ) and by Cter +viral RNA 10-fold higher (0.00408 ± 0.00143 clusters per µm^2^) (Table 3). In comparison, N cluster density is 0.0011 ± 0.00215 clusters per µm^2^ and in the presence of the viral RNA is 0.00152 ± 9.92e-4 clusters per µm^2^, showing that N is 3-fold less efficient in forming PI clusters and to multimerize on a membrane, suggesting that the N-ter domain could regulate the C-ter domain in binding lipids. These results strongly suggest a regulatory role of the N-ter on the C-ter for accessing anionic lipid binding sites, especially in the presence of the viral RNA. Most probably, the C-ter domain contains many lipid-binding sites. These lipid-binding sites may not all be accessible when N is full or when associated with the viral RNA.

**Table 3:** Size and Density of TF-PI clusters on simple PC:PS:PI SLB in the presence of the different N proteins (Full-length N, N-ter and C-ter proteins purified with His-tag), with or without viral RNA (vRNA). Cluster and density data are represented as the Median±IQR (Q3 upper quartile 75%-Q1 lower quartile 25%) of the number of clusters per frame issued from 3 independent experiments.

| Sample | Cluster Area |  | Cluster Density |  |
| --- | --- | --- | --- | --- |
| | Median ($\mu\text{m}^2$) | IQR (Q3-Q1) | Median ($\mu\text{m}^{-2}$) | IQR (Q3-Q1) |
| N ter only | 0.346 | 0.519 | 4.41E-04 | 1.65E-04 |
| N ter + RNA | 0.19 | 0.312 | 3.03E-04 | 3.86E-04 |
| C ter only | 0.606 | 1.219 | 7.72E-04 | 2.48E-04 |
| C ter + RNA | 0.779 | 1.488 | 0.00408 | 0.00143 |
| N only | 0.277 | 0.831 | 0.0011 | 0.00215 |
| N + RNA | 0.577 | 1.953 | 0.00152 | 9.92E-04 |

Furthermore, as reported above, in the case of N+vRNA, PI clustering adopted a circular structure (Figure 3E). In contrast, different structures were observed with the C-ter of N (Figure 4D). Instead, Cter+vRNA displayed two distinct morphologies on membranes: short filaments and dense PI-lipid–enriched clusters. These observations suggest that Cter-N assembles into a structure that is distinct from that formed by full-length N on lipid membranes.

### In vitro RNP assembly on biomimetic membranes recapitulates the size and volume of the viral core

Following the observation that N multimerizes and assembles with viral RNA on lipid membranes in vitro to form tight lipid–RNP interactions, we characterized the three- dimensional morphology of RNPs on both simple PC:PS:PI and ERGIC-like membranes. We first control the height of the lipid membrane alone in buffer solution after an incubation of 30 minutes at 37°C, as in our confocal measurement assays, on simple PC:PS:PI (Figure 5A) and ERGIC membranes (Figure 5B); we found a height of 1.63 ± 0.19 nm as expected for SLBs. Upon addition of the N protein alone on a PC:PS:PI SLB, we found objects with an average height of 3.79 ± 0.46 nm. The same experiments done on ERGIC membranes lead to the obtention of 8.26 ± 2.88 nm height objects (Figure 5C-D, G): this significant (2 -fold) increase in height shows the existence of bigger N complexes on the ERGIC-like membrane, suggesting an enhanced catalytic effect of the ERGIC lipids on N self-assembly. These heights of the N clusters match well with the previously reported size of the N proteins^29^. When adding RNA to N, we observed RNP objects of 13.01 ± 9.13 nm height on average in PC:PS:PI simple SLBs, while this average height reaches 38.92 ± 10.23 nm on ERGIC membranes (Figure 5E-F, H). The successive increases in height upon RNA addition to N and upon increased complexity of membrane lipid composition (ERGIC vs PC:PS:PI) nicely correlate with the previous fluorescent confocal microscopy observations (Figure 3, Table 1). Interestingly, the volumes of N/RNP assemblies obtained from the AFM measurements, comparing simple and ERGIC membranes, shows that RNP made on an ERGIC-like membrane reaches a size of 0.002 ± 0.00434 µm^3^ (Figure 5I), which is in the range of the average measured volume of a viral core^30–32^.

### Intracellular N clustering is enhanced by the presence of the viral RNA albeit no difference in VLP production

We further aimed at mimicking SARS-CoV-2 particle production in cells thanks to the transfection of plasmids expressing the structural proteins M, N and E in the absence or in the presence of the viral RNA. We evaluate the role of the viral RNA (same viral RNA as in vitro experiments containing the packageable signal recognised by N ^17^ in generating RNP clusters in cells and, if any, on Virus-Like Particle (VLP) production.

To get more insight into particle assembly and release processes, the production of Virus-like Particle (VLP) made of MNE was analysed in the presence and in the absence of the viral RNA in comparison to N alone or NE that can be released in the form of extracellular vesicles (EVs) (Figure 6 A-B). As expected, N expressed alone in cells or in the presence of the viral RNA did not release efficiently (Figure 6A, EVs, lanes 1 and 2). Since it has been described that the M/E protein is required for efficient assembly^6,7^. The presence of M, N and E, with or without viral RNA, was analysed and allowed efficient VLP production, leading to around 59%±10 of M release with vRNA and 57%±12 without vRNA (Figure 6B). This result indicates that there is no significant or no measurable difference in VLP production in the presence of vRNA by Western Blot. N incorporated into the VLP was estimated, but also showed no difference in the absence or in the presence of the vRNA (51%±13 versus 52%±8, respectively) (Figure 6B). In the absence of M, N+E is able to exit the cell with a good efficiency (33%±7) without any change with vRNA (Figure 6B). Although in the presence of M and E, VLP release remains the most efficient, with 2-fold more than the % of N+E vesicle release (compare 52%±8 to 33%±7, *p) (Figure 6B). Since the presence of vRNA did not change the overall VLP release, we decided to go further and analysed N clusters distribution in the cell cytoplasm. To visualise intracellular N, N-mEOS2 protein was expressed in pulmonary cells in the presence of M, N and E in proportion applied to produce VLP ^6^ in the presence or absence of vRNA (Figure 6C). We checked if a fluorescent tagged N introduction was forming VLP and if the presence of viral RNA was changing VLP release thanks to NTA measurement (Supplemental Figure 6): we could then compare VLP concentration and observe no measurable difference in the amount of VLP production with or without viral RNA, but could confirm that fluorescent MNE VLP are producing one-log more particles than NE or N containing vesicles (Supplemental Figure 6). Using TIRF-PALM on fixed pulmonary cells, we imaged intracellular cytoplasmic N clusters, labelled by N-mEOS2, and observed clusters’ locations in the cell cytoplasm (around 2/3) and at close proximity or on ERGIC-labelled membranes (around 1/3) (Figure 6C). We quantified and compared the sizes and numbers of N-mEOS2 labelled clusters in the presence or in the absence of viral RNA (Figure 6D). The results show that N/N-mEOS2/ME recruits N proteins into clusters more efficiently in the presence of viral RNA than in its absence. Although the median size of clusters did not appear to be significantly different (70+/- 19nm MNE, 72+/- 24 nm MNE+RNA, median+/- IQR), in the presence of RNA, clusters contain significantly more N localisations (Figure 6E-F). In addition, at low N density, N protein partitions into clusters in the presence of viral RNA, whereas they remain dispersed when viral RNA is absent, suggesting that viral RNA enhances the formation of RNP within the cell, at low N concentration (Figure 6G). The more N density, the more we observe the presence of N inside the clusters, suggesting an increase of N self-oligomerization (Figure 6G). This perfectly mirrors our in vitro data of RNP assembly observed in solution (Figure 1F) and on ERGIC membranes (Figures 3 and 4).

**Figure 6.**
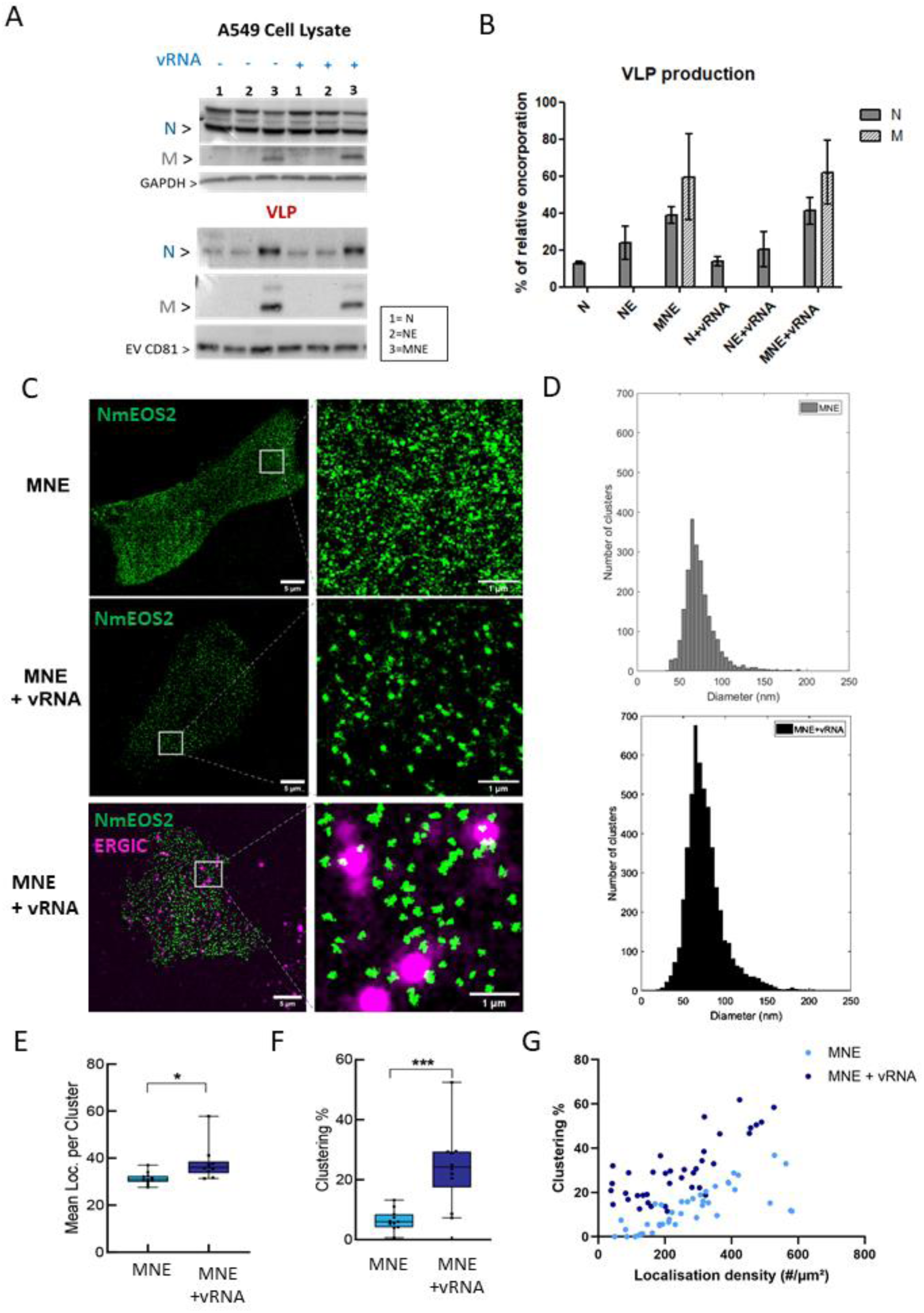
SARS-CoV-2 N clustering in the presence of the viral RNA in pulmonary cells producing Virus-like Particles. **A**) Western blot to evaluate the production of MNE Virus-like Particles (VLP) in the absence or the presence of viral RNA (vRNA) or N or NE containing extracellular vesicles (EVs) from transfected A549 pulmonary cells. **B**) Quantification of the % of N content in MNE VLP or N containing EVs in the absence or in the presence of viral RNA (vRNA) normalized to the total amount of N protein found in the cell lysates (CL) (normalized to GAPDH, the loading control). EVs production is estimated by the presence of the biomarker CD81. Data represents the Mean ± SD. N= 3 independent experiments. **C)** TIRF-PALM imaging of N-mEOS2 clusters in A549 pulmonary cells expressing MNE/N-mEOS2 in the presence or absence of viral RNA. ERGIC membranes are labelled using an anti-ERGIC53- AF647 antibody. The scale bar is 5µm for the main images and 1µm on the zoom images. **D**) Quantification of N-mEOS2 cluster size distribution in A549 cells with or without viral RNA. **E**) Each black dot represents an individual cell, and results are averaged per cell. Clusters were identified using a DBScan algorithm and their content is displayed as the mean number of localisations per clusters. **F**) The clustering efficiency (Clustering %) was calculated as the ratio of localisations within clusters to the total NmEOS2 localisations per cell (C). Data are presented as box-and-whisker plots, showing the median, 1st and 3rd quartiles, and minimum/maximum values. An average of 10 cells were analyzed per condition. Differences were evaluated using the Kolmogorov-Smirnov test (*p < 0.05, ***p< 0.001). **E**) Clustering efficiency was calculated for multiple ROIs (>3 per cell) and plotted as a function of localization density. Each dot represents one ROI.

These results suggest that, in eukaryotic cells, the presence of viral RNA promotes intracellular N protein clustering more efficiently, albeit with no significant apparent efficiency on overall VLP production.

## DISCUSSION

Our study provides mechanistic insight into how the SARS-CoV-2 nucleocapsid (N) protein engages with host membranes to drive ribonucleoprotein (RNP) assembly at the ER–Golgi intermediate compartment (ERGIC). Using biomimetic membrane systems, we show that full-length N displays low intrinsic affinity for negatively charged lipids such as phosphatidylinositol (PI) and phosphatidylserine (PS) (Figure 2), yet is capable of inducing pronounced lipid clustering (Figure 3). This apparent discrepancy suggests that membrane association is not governed solely by binding affinity, but rather emerges from cooperative multimerization of N on the membrane surface.

A key finding of this work is that viral RNA strongly enhances N-driven lipid reorganization. In the presence of viral RNA, N forms larger and more stable clusters, consistent with a model in which RNA acts as a scaffold or catalyst for N multimerization. This is supported by fluorescence correlation spectroscopy measurements indicating stronger N–lipid interactions under these conditions, particularly on ERGIC-like membranes. The reduced lipid diffusion within clusters further indicates that N multimerization reorganizes the membrane into dynamicaly caged/confined domains, potentially creating a favorable platform for RNP assembly.

The role of membrane composition is equally critical. ERGIC-mimicking membranes not only enhance lipid clustering but also promote the formation of RNP assemblies with dimensions comparable to those of the viral core (Figure 5). Atomic force microscopy (AFM) reveals an increased height in N-protein clusters on ERGIC membranes compared to PCPSPI membranes (Figure 5). This observation, supported by the flexible conformational ensemble and dimeric state characterized by SAXS data and EROS modelisation, suggests a membrane-dependent mechanism for nucleocapsid multimerization^29^ . It also reveals that RNP complexes formed on these membranes reach volumes on the order of 0.002 µm³, consistent with reported viral core sizes^30,31^. In contrast, assemblies formed on simpler membranes remain significantly smaller, highlighting a synergistic interplay between lipid environment and RNP organization. These observations are in agreement with lipidomic analyses showing enrichment of PI in SARS- CoV-2 particles^33,34^, supporting the idea that PI-rich ERGIC membranes provide a privileged platform for early viral assembly. Later assembly steps may require the adjunction of other type of lipids, like M specific binding to the Golgi-enriched anionic lipid Ceramide-1- phosphate that is required for S/E recruitment into the viral particle^35^.

The C-terminal domain of N retains the ability to bind membranes and induce lipid clustering, whereas the N-terminal domain alone is insufficient (Figure 4), in agreement with a previous study^16^. Interestingly, the C-terminal domain generates a higher number of lipid clusters than the full-length protein, albeit with altered density and organization, suggesting a more aspecific lipid binding to PS/PI lipids. This can also suggest that the N-terminal domain may act as a regulatory module, potentially modulating N interactions with both viral RNA and more specific lipids. One possibility is that the N-terminal domain, which is known to bind viral RNA^14^, constrains or spatially organizes N multimerization, preventing uncontrolled lipid clustering and core size formation. In this context, viral RNA binding may relieve this regulation, promoting coordinated assembly of lipid-associated RNPs. These aspects would require further investigations.

The morphology of N–lipid assemblies also provide important clues. We observe diverse structures, including dense clusters, hollow or ring-like assemblies, and filamentous organization, particularly in the presence of RNA (Figure 3E & 4). The emergence of ring-like structures suggests that N may bridge viral RNA into circular or looped conformations, which could facilitate compaction of the long viral genome during packaging^36^. Differences between simple PC:PS:PI and ERGIC-like membranes further indicate that lipid composition influences not only the extent but also the geometry of RNP assembly.

Although our in vitro system captures key features of N–lipid–RNA interactions, it also has limitations. Notably, the N protein used here is not phosphorylated, whereas phosphorylation of the serine/arginine-rich linker region is known to regulate N function^37,38^. This post-translational modification likely modulates the balance between RNA binding, protein multimerization, and membrane association, and may be critical for switching between roles in replication and assembly. Future work using phosphomimetic mutants will be necessary to assess how phosphorylation impacts lipid-dependent RNP formation.

Another important limitation is the absence of other viral structural proteins, particularly the membrane (M) protein. M is thought to play a central role in virion assembly and has been proposed to interact with N and RNP complexes, although direct evidence remains limited^20^ . Our results suggest that lipid-associated RNPs could serve as intermediates for recruitment to ERGIC membranes containing M, E, and S proteins. Reconstituting such systems with transmembrane M protein in supported lipid bilayers would be a major experimental advance, albeit technically challenging.

Finally, our cellular data indicate that viral RNA enhances N clustering in vivo, consistent with our in vitro observations, yet without detectable changes in virus-like particle production using immunoblots or fluorescent particle counting by NTA (Figure 6). This suggests that while RNA-mediated N clustering is important for organizing RNPs, additional factors likely governlater steps of virion formation and release. Indeed, the experiments are optimized for detecting VLPs in cell supernatants via high, constitutive expression of the proteins. Therefore, the physical availability of membranes for assembly, cellular enzymes involved in particle maturation (e.g., glycosylation), and the dynamics of the secretion mechanism itself may act as bottlenecks, limiting production to a level where differences cannot be observed when vRNA is assessed. Further experiments using low-expression systems and appropriate detection methods to shift the bottleneck to the assembly step might yield different results.

In conclusion, our findings support a model in which cooperative interactions between N, viral RNA, and PI-rich ERGIC membranes drive the formation of lipid-associated RNP assemblies with dimensions and properties consistent with the viral core. This lipid-dependent step likely occurs early during assembly, following genome export from replication compartments, and may represent a critical intermediate in virion formation. These findings position lipid–protein coupling as a central determinant of early coronavirus assembly and provide a framework for dissecting how membrane composition shapes viral morphogenesis.

## MATERIAL AND METHODS

### Plasmids cloning

The plasmids expressing the SARS-CoV-2 (Wuhan) N, M, E and N-GFP proteins in eukaryotic mammalian cells were previously described^6^. The plasmid pET30a-SARS-CoV2-NP expressing the SARS-CoV-2 (Wuhan) N protein in procaryotic (bacterial) cells was obtained from Addgene (# 165090). The plasmid encoding the viral RNA was generated by Gibson Assembly using the NEBuilder® HiFi DNA Assembly Master Mix (New England Biolabs, E2621S). PCR fragments corresponding to the 5’ untranslated region (5’UTR) of SARS-CoV- 2, synthetic tag in-frame with the start codon, and the 3’UTR of SARS-CoV-2 were assembled into a pcDNA backbone. All viral sequences were derived from the original Wuhan reference strain (NC_045512.2). The Nucleocapsid N-ter (1-212) and C-ter (213-422) expressing plasmids were generated using NEBuilder HiFi DNA Assembly Master Mix with the primers N N-ter and C-ter listed (Supplemental Table 1) and inserted into the pET30a-SARS-CoV2- NP digested with HindIII /BamHI.

**Supplemental Table 1:**
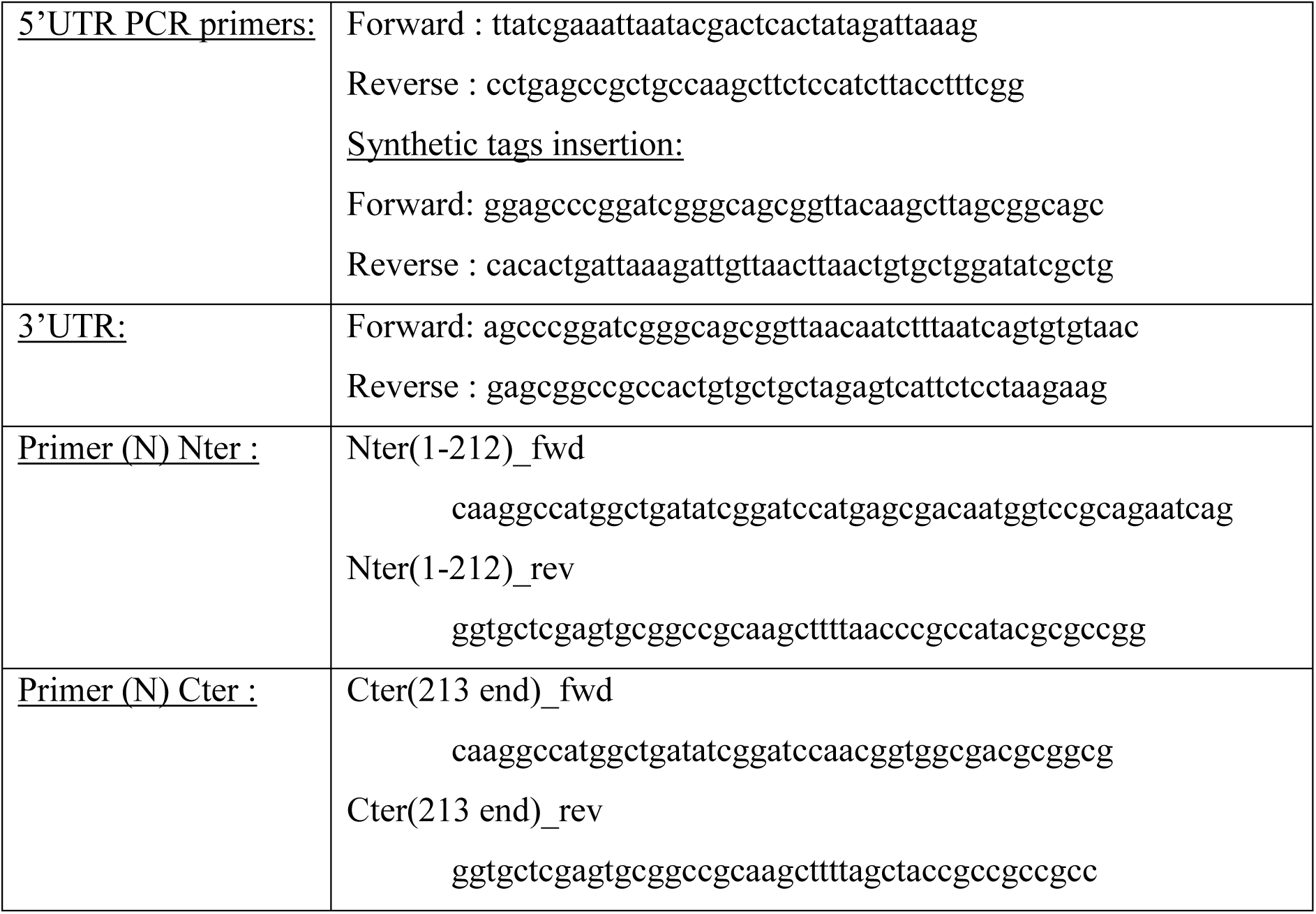
The primers sequences used for DNA cloning.

### Expression and purification of the SARS-CoV-2 N proteins from bacteria

For preparation of chemically competent bacteria, an overnight bacterial culture was inoculated into LB broth at a dilution of 1:100 and incubated with shaking at 200 rpm at 37°C until the culture reached mid-log phase (OD_600_ = 0.45-0.55). Following a 15 min incubation on ice, the bacterial culture was centrifuged at 3000g at 4°C for 10 minutes. The supernatant was discarded and the bacteria were resuspended in 0.1 volumes (compared to the original culture volume) of ice-cold RF1 buffer (100 mM RbCl, 50 mM MnCl_2_, 30 mM potassium acetate, 10 mM CaCl_2_, 15% (v/v) glycerol, pH 5.8). Following 10 min of incubation on ice, the bacteria were pelleted by centrifugation as described above. The supernatant was discarded and the bacterial pellet was resuspended in 0.01 volumes (compared to the original culture volume) of ice-cold RF2 buffer (10 mM MOPS, 10 mM RbCl, 75 mM CaCl2, 15% (v/v) glycerol, pH 6.5). 200 µl aliquots were snap-frozen in a dry ice/ethanol bath and stored at -80°C. For each heat shock transformation, 50 µl of competent bacteria were thawed on ice before addition of 5 µg of plasmid DNA. Bacteria were then incubated for 30 min on ice before being heat shocked for 1 min at 42°C. Following addition of 1ml of LB broth, the transformation mix was then incubated shaking at 37°C with shaking at 200 rpm for 1 h. Bacteria were then plated on LB agar plates containing the appropriate antibiotic and incubated at 37°C overnight.

#### #Purification of SARS-CoV-2 N protein

An overnight (O.N.) pre-culture was grown at 37°C and used to inoculate four large flasks containing three liters of LB culture supplemented with Kanamycin (50 μg.mL^−1^). The culture was grown until the optical density at 600 nm reached 0.8 and was plunged into a melting ice bucket for 45 min to enhance production of chaperones. Protein expression was then induced by adding 1 mM of Isopropyl β-D-1-thiogalactopyranoside (IPTG) and the culture was incubated O.N. at 18℃ under agitation. The day after bacteria were pelleted by a 15 min centrifugation step at 6000 *g*. The pellet was resuspended in lysis buffer: 50 mM Tris-HCl pH 8, 500 mM NaCl, 10 mM Imidazole, 1 mM Benzamidine and one tablet of EDTA-free Protease Inhibitor Cocktail (Roche). Cell lysate was sonicated 4 times during 2 min (2 sec pulse + 2 sec pause) at 40 % intensity (Digital Sonifier, Branson) followed by a 45 min centrifugation step at 27000 *g*. The supernatant was loaded twice onto 4 gravity columns each containing 3 mL of Ni-NTA Sepharose beads. The beads were washed with 10 column volume (CV) of lysis buffer. The column was washed on more time with 10 CV of 50 mM Tris-HCl pH 8, 500 mM NaCl, 40 mM Imidazole and 1 mM Benzamidine. The protein was eluted with 3 CV of 50 mM Tris-HCl pH 8, 500 mM NaCl, 200 mM Imidazole and 1 mM Benzamidine. The eluted protein was dialyzed O.N. at 4℃ against 50 mM Tris-HCl pH 8 and 500 mM NaCl. The protein was concentrated to 5mg.mL^−1^ before being injected onto a Superdex 200 Increase 10/300 GL column and eluted in 50 mM Tris-HCl pH 8 and 500 mM NaCl. The pure fractions were concentrated on a centricon with a 10 kDa cut-off to 35 μM, aliquoted, flash-frozen in liquid nitrogen and stored -80°C.

#### #Purification of tag-free SARS-CoV-2 N protein

The protein was expressed and purified similarly as for the tagged version of SARS-CoV-2 N protein. After elution of the Ni-NTA Sepharose, thrombin was added to the protein (4U of thrombin per mg of SARS-CoV-2 N protein) and the mix was dialyzed against 50 mM Tris-HCl pH 8 and 500 mM NaCl O.N. at 4℃. The protein was then passed three times through a 2 mL Ni-NTA Sepharose column equilibrated with the same buffer as the dialysis one. The protein collected in the flow-through was concentrated to 3 mg.mL^−1^ and further purified as described for the tagged version on size-exclusion chromatography and eluted in 50 mM Tris-HCl pH 8 and 500 mM NaCl. The peak fractions were pooled, concentrated to 29 μM, aliquoted, flash-frozen in liquid nitrogen and stored -80°C. (see Supplemental Figure 1A-B).

#### #Purification of SARS-CoV-2 N-terminal and C-terminal domains

The N-terminal and C-terminal domains of the SARS-CoV-2 N protein were essentially purified with the same purification procedure as for the full-length protein to the exception that the culture volume was 6 liters and that the size-exclusion chromatography step was performed on a Superdex 75 Increase 10/300 GL column. The proteins were stored at a final concentration of 23 μM or 30µM (depending on the batch). (Supplemental Figure 1 C-D-E).

### In vitro transcribed and purified viral RNA

The plasmid containing the viral RNA sequence was linearized prior to transcription. The plasmid was digested by NotI-HF and MluI-HF (New England Biolabs, R3189S and R3198S). In vitro transcription was performed using the HiScribe™ T7 ARCA mRNA Kit (New England Biolabs, E2065S) to generate capped and polyadenylated RNAs. Transcribed RNAs were purified using the RNeasy Mini Kit (Qiagen, 74104) and stored at –80°C. RNA integrity was verified by agarose gel electrophoresis (Figure 1D). The uncapped and unpolyadenylated RNA length was 2428 bp. This in vitro transcribed RNA was named *viral RNA* and its concentration used in this study was 0.2 nM. The Molecular Weight (MW) of the viral RNA is 777 kDa for a lenght of 2425 bases.

### N/RNP size measurement in solution using the Videodrop®

The Videodrop instrument (Myriade) has been used to estimate the size of the SARS-CoV-2 purified nucleocapsid N, N-ter or C-ter proteins [1µM] and the RNP made of N proteins in the presence and absence of the viral RNA [0.2nM] in solution [buffer: Tris (150 mM NaCl, 10 mM Tris– HCl, pH 7.4)]. This instrument can measure the size of particles between 80 nm to 500 nm diameter range for a concentration below 8.10e5 particle/mL.

### Large Unilamellar Vesicles (LUVs) preparation

Extracted Lipids used in this study were received from Avanti Polar Lipids. Lipid stocks were prepared in chloroform:methanol (2:1) mixture. The required volume of lipid was taken in a round-bottom flask and evaporated using a vacuum pump coupled with a rotavapor to form lipid films. lipid films were hydrated in Tris-buffer (150 mM NaCl, 10 mM Tris–HCl, pH 7.4) using freeze-thaw cycles and extruded through a 100/200-nm Whatman polycarbonate filter (GE Healthcare). The size of the LUVs was confirmed using dynamic light scattering (DLS) on a Malvern analytical Zetasizer Nano (Figure 2). LUVs used in the study have been tabulated in table 3. The concentration of lipids used was 0.2mg/ml.

**Supplemental Table 2:** The lipid compositions of model (LUV or SLB) membranes used in this study.

| Lipid systems | Compositions |
| --- | --- |
| POPC | 100:0 |
| PC:PI | 80:20 |
| PC:PS | 80:20 |
| PC:PS:PI | 80:10:10 |
| PC:PI | 90:10 |
| PC:PI(3)P | 95:5 |
| PC:PI(4)P | 95:5 |
| PC:PI(3,4)P2 | 97.5:2.5 |
| PC:PI(3,5)P2 | 97.5:2.5 |
| PC:PI(4,5)P2 | 97.5:2.5 |
| PC:PI(3,4,5)P3 | 98.34:1.66 |
| PC:PS:PI(3)P | 85:10:5 |
| PC:PS:PI(4)P | 85:10:5 |
| PC:PS:PI(3,4)P2 | 87.5:10:2.5 |
| PC:PS:PI(3,5)P2 | 87.5:10:2.5 |
| PC:PS:PI(4,5)P2 | 87.5:10:2.5 |
| PC:PS:PI(3,4,5)P3 | 88.34:10:1.66 |
| ERGIC-like<br>PC:PS:PI:PE:Chol | 50:10:10:20:10 |

### In vitro LUV co-sedimentation assays

SARS-CoV-2 N protein (2 µM) was incubated with LUVs of 200 nm diameter (1 mg/ml) in a final volume of 100μl at room temperature for 30 min. Samples were then centrifuged at 220,000g in a Beckman TLA 100 rotor at 4 °C for 30 min. Each sample was then divided into supernatant (S = 80 μl), containing unbound protein, and pellet (p = 20 μl), containing LUV-bound protein. P was diluted in 20 μl of Tris–NaCl buffer (150 mM NaCl, 10 mM Tris–HCl, pH 7.4) to maintain the equivalence between the S and P volumes. Twenty microliters of S and P were analysed by SDS-PAGE and protein were detected by staining with Coomassie Blue. The protein intensities (intensity of supernatant: Is, intensity of pellet: Ip) were quantified using the Fiji ImageJ software. The percentage of LUV-bound protein was calculated as: Percentage of protein present in the pellets = (I_P_/I_P_+I_S_) *100

I_P_: Intensity of the pellets band

I_s:_ Intensity of the supernatant band

### In vitro LUV-N float-up assays

N-protein affinity for membranes was measured by performing float-up experiments in which N-protein was mixed with large unilamellar vesicles (LUVs), and the N-protein–bound fraction was separated from the free fraction using a density gradient. LUVs were prepared by drying a lipid mixture of the chosen composition (generic composition PC : anionic lipid : C18 DiI at a 79.5 : 20 : 0.5 mol% ratio) dissolved in chloroform under vacuum. The lipid films were then rehydrated in buffer (150 mM NaCl, 10 mM Tris, pH 7.4) to reach a final concentration of 5.5 mM. Rehydrated lipid solutions were then frozen and thawed 10 times before being extruded 21 times through a 100-nm polycarbonate filter (Avanti Research). LUVs were then mixed with N-protein to reach a final lipid concentration of 5 mM and a theoretical N-protein concentration of 3 µM. The protein–LUV solution was incubated at room temperature for 30 minutes to allow equilibration.

Following incubation, the LUV–protein solution was mixed with OptiPrep density gradient medium (CAS n°92339-11-2) to obtain a final OptiPrep concentration of 30% (w/v). Density gradients were then formed in Beckman-Coulter polycarbonate 8 × 34 mm ultracentrifuge tubes by layering 200 µL of the LUV–protein solution, 300 µL of 20% OptiPrep in NaCl buffer, and finally 150 µL of buffer. Tubes were ultracentrifuged at 250,000 g, 4°C for 3 hours using an Optima MAX-XP Ultracentrifuge equipped with a TLA120.1 rotor (Beckman-Coulter). After ultracentrifugation, the floated LUVs were collected by pipetting 100 µL from the top of the tubes. Due to the presence of 0.5 mol% C18 DiI lipids, the LUV fraction could be visually confirmed as fully collected. To ensure sufficient signal on gels, collected fractions were concentrated by evaporating water at 70°C, resulting in a 2- to 3-fold concentration of the floated fraction.

After collection, the N-protein content in the floated fraction and the initial protein concentration before flotation were quantified by PAGE gel electrophoresis on a 10% acrylamide gel. Protein concentration was measured on the gel using a BSA titration standard curve, providing the bound protein concentration [P]_bound_ and the total protein concentration [P]_total_. From these concentrations, we estimated the dissociation constant Kd of the N-protein toward a membrane of defined composition, with K_d_ defined as:

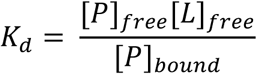

Considering the large excess of lipid concentration compare to the protein concentration even if all protein were bound to the membrane, we have [*L*]*_free_* ≈ [*L*]*_total_*. Thus, K_d_ can be written as :

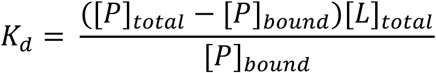

From the gel quantification, we observe the bound N-protein concentration to be around or even below 1µM, meaning the anionic lipids are present in a roughly 1000-range excess relative to N-protein.

### SLB preparations

Two methods were used. Vesicle fusion method for simple SLBs preparation was used for SLB made of PC:PI or PC:PS or PC:PS:PI. The cover slips were cleaned with piranha solution (sulfuric acid (H2SO4): hydrogen peroxide (H2O2) 3:1)/ozone treatment for 30 min. LUVs with desired lipid compositions and with fluorescent lipid (TMR-PI or TopFluor-PS) in Tris buffer (150 mM NaCl, 10 mM Tris– HCl, pH 7.4) were prepared according to the method describe above. 0.2mg/ml of LUVs in the cleaned glass cover slips were deposited and kept at 37°C for 30 min for SLBs preparation. The coverslips were washed with Tris-buffer to clean unfused vesicles. PCPSPI and ERGIC SLB preparation used Langmuir Blodgett (LB)^23^ : The lipid bilayer with composition PC:PS: PI (80:10:10) or ERGIC mimicking membrane and with fluorescent lipid (TF-PI) was prepared using LB techniques. The process was conducted in a Langmuir trough with a deionized water subphase maintained at 20 °C. The trough temperature was maintained at 20◦C using a Julabo chiller (purchased from Srico Pvt. Ltd.), with water circulation beneath the trough through a metal body. A glass substrate, rendered hydrophilic via RCA cleaning, was submerged beneath the water surface of the trough and secured using a dipper controlled by KSV Nima software. An aliquot of the lipid mixture was spread on the water surface to achieve a surface pressure of 3–5 mN/m. The bilayer was subsequently deposited onto the substrate via a sequential upstroke at 5 mm/min, followed by a 400-second drying interval, and a final downstroke at 3 mm/min. The completed bilayer was then stored in an aqueous medium and a through wash with Tris (150 mM NaCl, 10 mM Tris– HCl, pH 7.4) buffer to maintain its stability.

Desired protein concentrations (1 µM) were directly added to the surface of SLBs (pre-incubated or not with 0.2 nM of viral RNA) and incubated for 30 min at RT for each condition. The RNP were formed by incubation of 1µM N with 0.2nM viral RNA and incubate at 37°C for 30 min before adding on to the surface of the bilayer lipidic membrane (SLB).

### Confocal Microscopy

Confocal images were generated using a LSM980-laser-scanning microscope (Zeiss) equipped with a 63×, 1.4 NA oil objective at the MRI facility, CNRS Montpellier, France. At the Physics Department, IISc Bangalore, India, optical microscopy images were acquired on a Leica SP5 microscope integrated with a Stedycon optical setup (Abberior Instruments GmbH). The same system was used for confocal Fluorescence Correlation Spectroscopy (FCS), a Picoharp 300 module and an LSM upgrade toolkit were used, with data acquisition managed by SymPho Time 64 software. All the images were processed with ImageJ/Fiji software. Subsequent image processing and quantitative cluster analysis were conducted using ImageJ/Fiji over at least 500 clusters to maintain high statistical confidence. FCS analysis was performed in QuickFit software (Langowski Group (B040), DKFZ) using a 2D diffusion model.

### Atomic Force Microscopy

Atomic Force Microscopy (AFM) was performed on lipid bilayers deposited onto glass coverslips. The bilayers were incubated with 1 µM N Protein, both in the absence and presence of 0.2 nM viral RNA. Following incubation, samples were extensively washed with Tris-buffer to eliminate unbound species from the bulk solution. The liquid cell was next filled with 2 mL Tris-buffer. The first set of measurements took place in the Physics Department at IISc Bangalore using a Park Systems NX-10 AFM. For this, an HQ:NSC14/No_Al-15 cantilever (no aluminum coating) was operated in non-contact tapping mode within a liquid cell filled with Tris-buffer. The cantilever was tuned to a resonant frequency of approximately 50–70 kHz in liquid (referenced from 160–170 kHz in air), with key imaging parameters set to a drive amplitude of 20 nm, a set point of ∼5 nm, a scan rate of 0.6–0.7 Hz, and an integral gain of ∼1. Additionally, a second set of experiments was performed at CEMIPAI, CNRS, Montpellier, France. The Bio-AFM imaging was performed on glass-bottom FluoroDish Cell Culture dishes (WPI) that were coated, or not, with PC:PS:PI (80:10:10) SLBs at room temperature on a NanoWizard IV atomic force microscope (JPK BioAFM, Bruker Nano GmbH, Berlin, Germany) mounted on an inverted microscope (Nikon Ti-U, Nikon Instruments Europe B.V, Amsterdam, the Netherlands) equipped with a standard monochrome CCD camera (ProgRes MFCool, Jenoptik, Jena, Germany).

### Cell lines and culture conditions

Pulmonary A549 adenocarcinoma human alveolar basal epithelial cell lines were obtained from ECACC (#86012804, Sigma-Aldrich, Germany) and cultured in Roswell Park Memorial Institute medium (RPMI) from Gibco supplemented with 10% heat-inactivated fetal calf serum (FCS, Thermo Fisher, USA), 50 U/mL of penicillin (Ozyme, France), 50 µg/mL of streptomycin (Ozyme, France), 1 mM sodium-pyruvate (Ozyme, France) and 25 mM HEPES (Ozyme, France), at 37 °C with 5% CO_2._

### Transfection and VLP/EV production

Cells were seeded into a 35mm Cell Culture dishes 24 h before transfection. Plasmids expressing M, N, and E proteins and a plasmid expressing the viral RNA of SARS-CoV-2 with a plasmid ratio of 3:12:2:12 respectively as determined previously ^6^, or M, N, E, N-GFP/N- mEos2 and a plasmid expressing the viral RNA, with a plasmid ratio of 3:9:2:3:12 respectively were transfected with Lipofectamine 3000 (Invitrogen) according to manufacturer’s protocol. A total of 2.2 µg of mixed plasmids per well was applied for the transfection. Cell lysate was collected by lysis into the RIPA buffer. Cell supernatant was collected and MNE-VLP (or N- EVs) were purified by ultracentrifugation against a 20% sucrose cushion in TNE buffer, pH7.4 at 130.000g for 2h at 4°C. The VLP/EV were then resuspended in sterile TNE with loading dye and deposited on a 10% SDS-PAGE and analysed by immunoblots to analyse the protein content of the VLP or EV. VLP production was calculated by quantification of the western blot thanks to ImageJ plugin, as % of VLP = VLP/[VLP + CL relative to GAPDH] x100.

### Measurement of VLP/EV particle concentration and size using a Nanoparticle Tracker Analyser

The analysis was performed using a ZetaView x30 analyzer (Particle Metrix), pre-calibrated with 100 nm polystyrene beads (Particle Metrix) to optimize focus. Purified VLP/EV particles were diluted in 10% PBS to ensure an adequate particle count for individual measurement. Specifically, MNE and MNE + vRNA particles were diluted at a 1:250 ratio, while NE, N, NE + vRNA, and N + vRNA particles were diluted at a 1:50 ratio. The ZetaView system was primed with 10 mL of 10% PBS, followed by the injection of 0.7 mL of the diluted particle suspension. For each sample, 3-4 measurements were conducted using a 488 nm laser and a 500 nm longpass filter, capturing 10 fields of view per measurement. Between each measurement, an additional 50–100 µL of the sample was injected. A second injection of freshly diluted particles was then processed identically. 30mL of water followed by 10mL of PBS were injected between each condition. The acquired measurements were analyzed using the manufacturer’s software (PEX, Particle Metrix), which provided size and concentration peak data for each field of view, taking into account the dilution.

### PALM TIRF Microscopy

A549 cells were seeded into a 35mm Glass-bottom FluoroDish Cell Culture dishes 24 h before transfection. Plasmids expressing viral M, N, NmEOS2 and E proteins were transfected with Lipofectamine 3000 (Invitrogen) according to manufacturer’s protocol. A total of 1.4 µg of mixed plasmids per well was applied for the transfection. Co-transfection of several plasmids was conducted with M, N, N-mEOS2, E and a plasmid expressing the viral RNA of SARS- CoV-2 with a plasmid ratio of 3:6:6:2:6 respectively as determined previously^6^. Then, transfected cells were washed 4 h later with phosphate-buffered saline (PBS 1x) and harvested 24 h post-transfection. The cells were then fixed with 4% PFA, wash and imaged in PBS for N-mEOS2 by photoactivable localization microscopy (PALM) coupled to TIRF-M imaging. For ERGIC labelling, cells were fixed with 4% PFA + 4% sucrose, washed three times in PBS and permeabilized with 0.1% Triton X-100 for 5 min at room temperature. Following, cells were blocked with 10% BSA for 30 min, then incubated with an anti-ERGIC53 antibody (C-6, sc-365158 AF647; Santa Cruz Biotechnology) in 10% BSA for 90 min at room temperature. Finally, cells were washed three times in PBS before observation. PALM images were generated using a TIRF microscope (Abbelight, France) equipped with a 100X, 1.49 NA Apo TIRF objective. Images were processed accordingly to the Abbelight manufacturer. Cluster numbering was determined thanks to Neo Analysis v40.4 software (Abbelight).

### Statistical tests

Data were analyzed using Origin and GraphPad/Prism softwares and statistical Mann-Whitney, Kolmogorov-Smirnov or t tests were applied to compare the different values or group of values, as indicated in the figure legends. In the main text, unless specified, data are presented as Median value ± IQR, interquartile range showing the difference between the 75th and 25th percentiles of the data.

## Data Availability

All relevant data supporting the key findings of this study are available within the article and in Supplementary Information files. Source data will be provided as a Source Data file in a ZIP folder containing the data of each graph and in Supplementary information upon request. Due to size constrains, relevant raw data of FCS are provided and available upon request, the remaining raw data are available upon reasonable request.

## Acknowledgments

We are grateful to CEMIPAI UAR3725 CNRS/University of Montpellier, France for supporting BSL3 facility and BSL3 Microscopy and for excellent technical services. We thank the Montpellier Microscopy Center (MRI) in France for confocal microscopy facilities and the GDR ImaBio for continuous support.

## AUTHOR CONTRIBUTIONS

JM, SS, JS, PM, LB, VR, JF, DA, SL and EM performed the experimental studies; JM, SS, JS, PM, VR, LB, MB and CF carried out the analysis. JB, CF and DM supervised the work. DM wrote the original draft. SS, JD, MB, CF, JB and DM reviewed and edited the manuscript. JB, CF and DM Funding acquisition. JB and DM project administration.

