## Supplementary material for "SARS-CoV-2 nucleocapsid protein engages with viral RNA and ERGIC lipids to drive viral core assembly": (Supplemental Figure

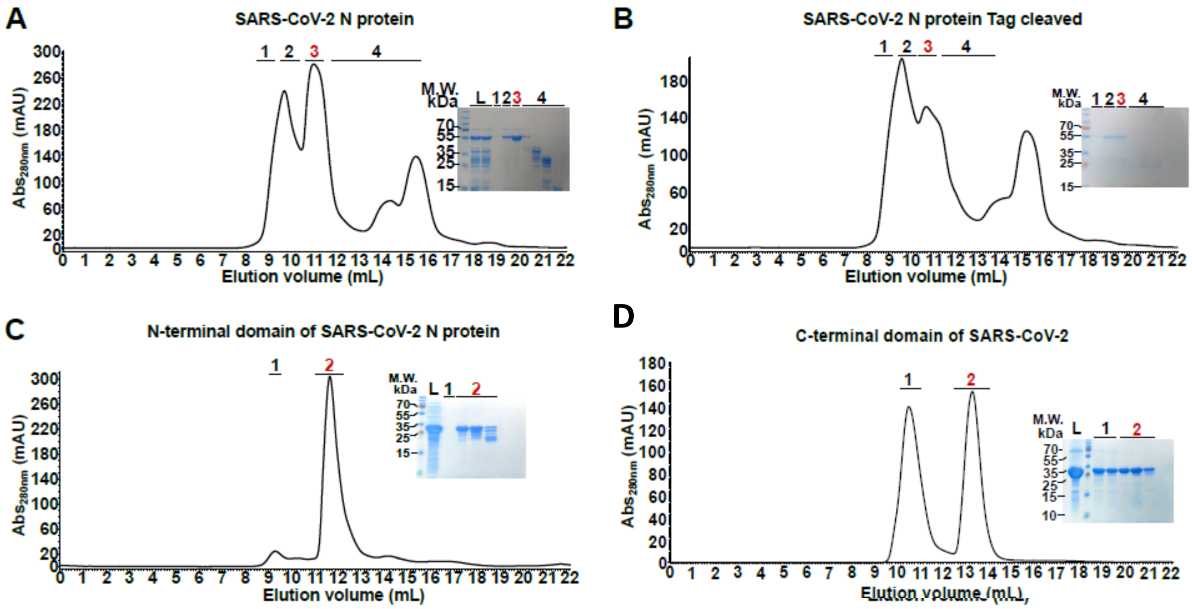


**Supplementary Fig. 1: Purification of the SARS-CoV-2 N protein wild-type and variant forms**

All the panels display the last purification step on size-exclusion chromatography for each variants: **A-**Full-length SARS-CoV-2 Wuhan N protein, **B-**Full-length SARS-CoV-2 Wuhan N protein with His tag removed, **C-**N-terminal domain of SARS-CoV-2 N protein, **D-**C-terminal domain of SARS-CoV-2 N protein. The red number and the line about each chromatogram indicate the peak fractions that were pooled and used for subsequent experiments. The purity of the proteins can be assessed on the corresponding Coomassie stained SDS-PAGE. L indicates the fraction loaded on the SEC column (meaning protein before the last polishing step) and T, on panel H-, indicates the protein before its tag being cleaved with the thrombin.

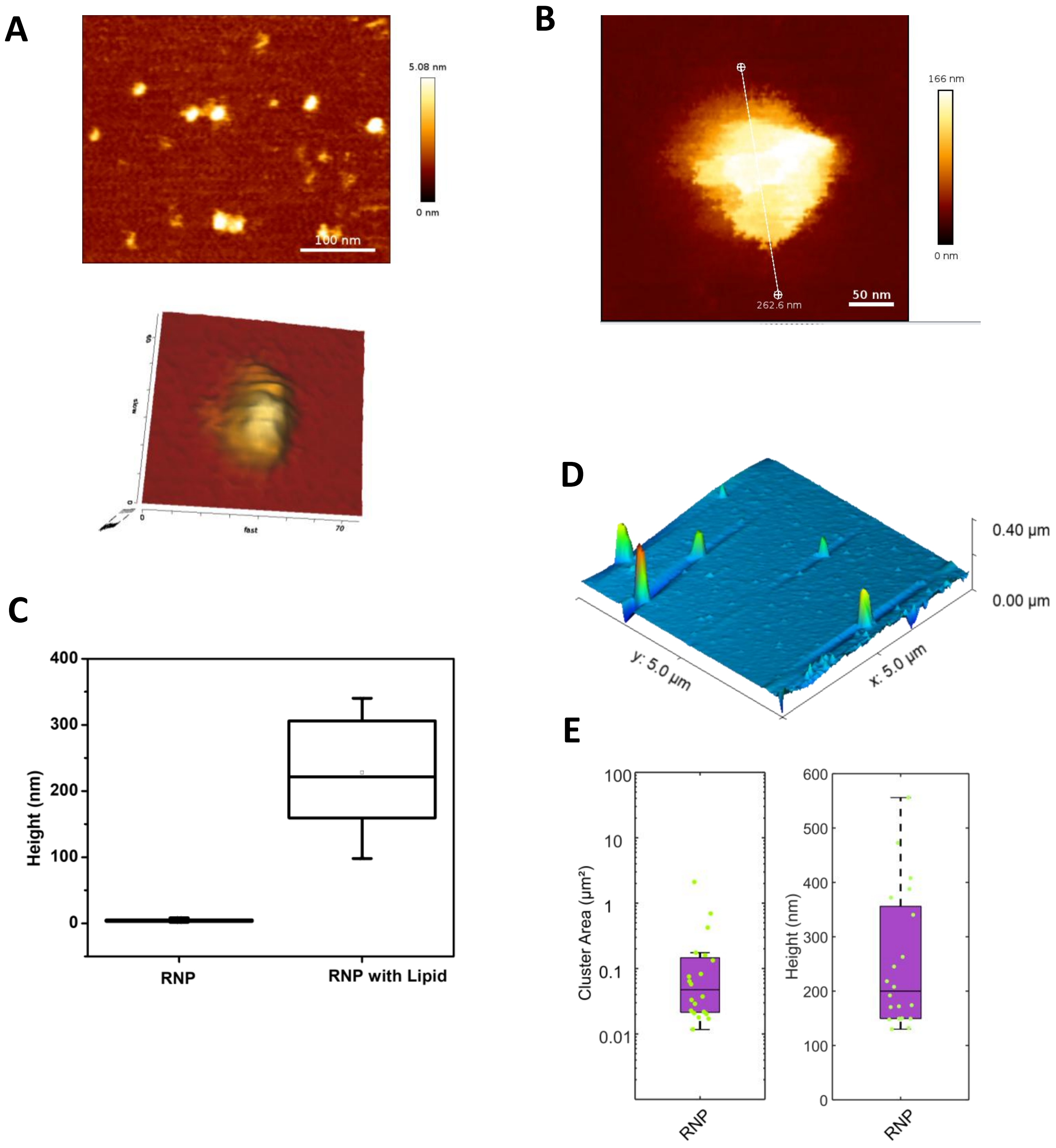


**Supplementary Fig. 2: Bio-AFM imaging of RNP deposit on glass surface versus on model membranes.** SARS-CoV-2 N and the viral RNA were incubated for 30min at 37°C in Buffer solution and deposited either on (**A**) on a glass coverslip or (**B**) on a simple PC:PI biomembrane for 2D/3D imaging using atomic force microscopy (Bio-AFM). Scale bar is 50nm. (**C**) Graph indicates the height of the RNP measured by Bio-AFM tapping mode comparing a deposit on glass surface versus SLB. (**D**) 3D AFM images of the RNP deposit on SLB. (**E**) Quantification of the RNP cluster area and height of the RNP on the SLB surface after 30 min deposit and washes.

A

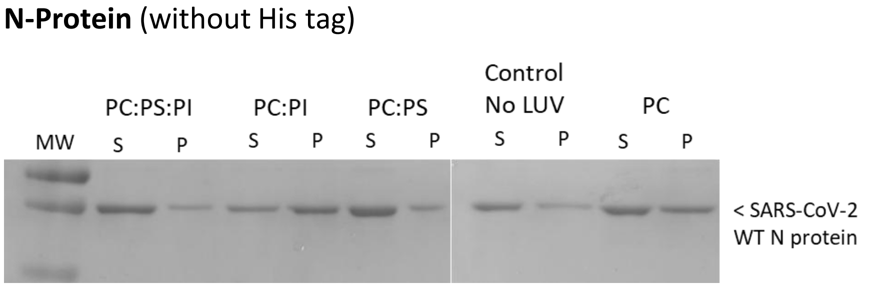


B

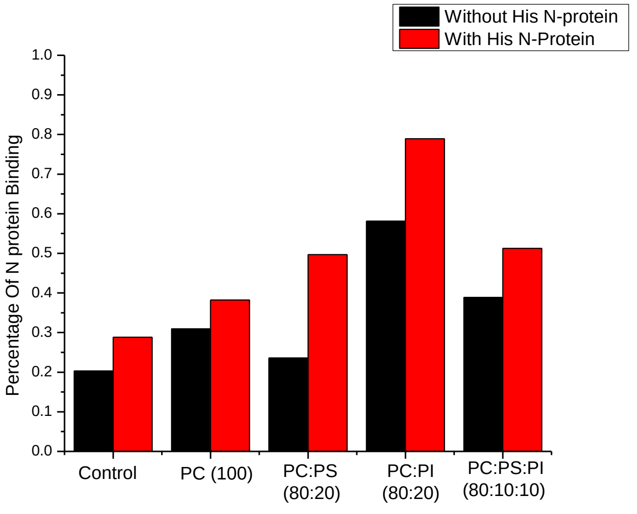


**Supplementary Fig. 3: LUV binding assay comparing N proteins with and without His tag.** (**A**) Example of Western blots of WT N protein without His tag in interaction with LUV of simple composition mixing PC, PS or PI lipid at different ratio, as indicated. (**B**) Comparative quantification of the % of N protein binding to the LUV Black color bar are the N protein without His tag and red color bar are N with His tag; % P=P/(P+S)x100; S = N protein in the Supernatant and P = N protein in the Pellet with LUV after centrifugation. All the Western Blot are revealed with a SARS-CoV-2 anti-N protein. N=2 independent experiments. P: pellet fraction; S: supernatant fraction after centrifugation.

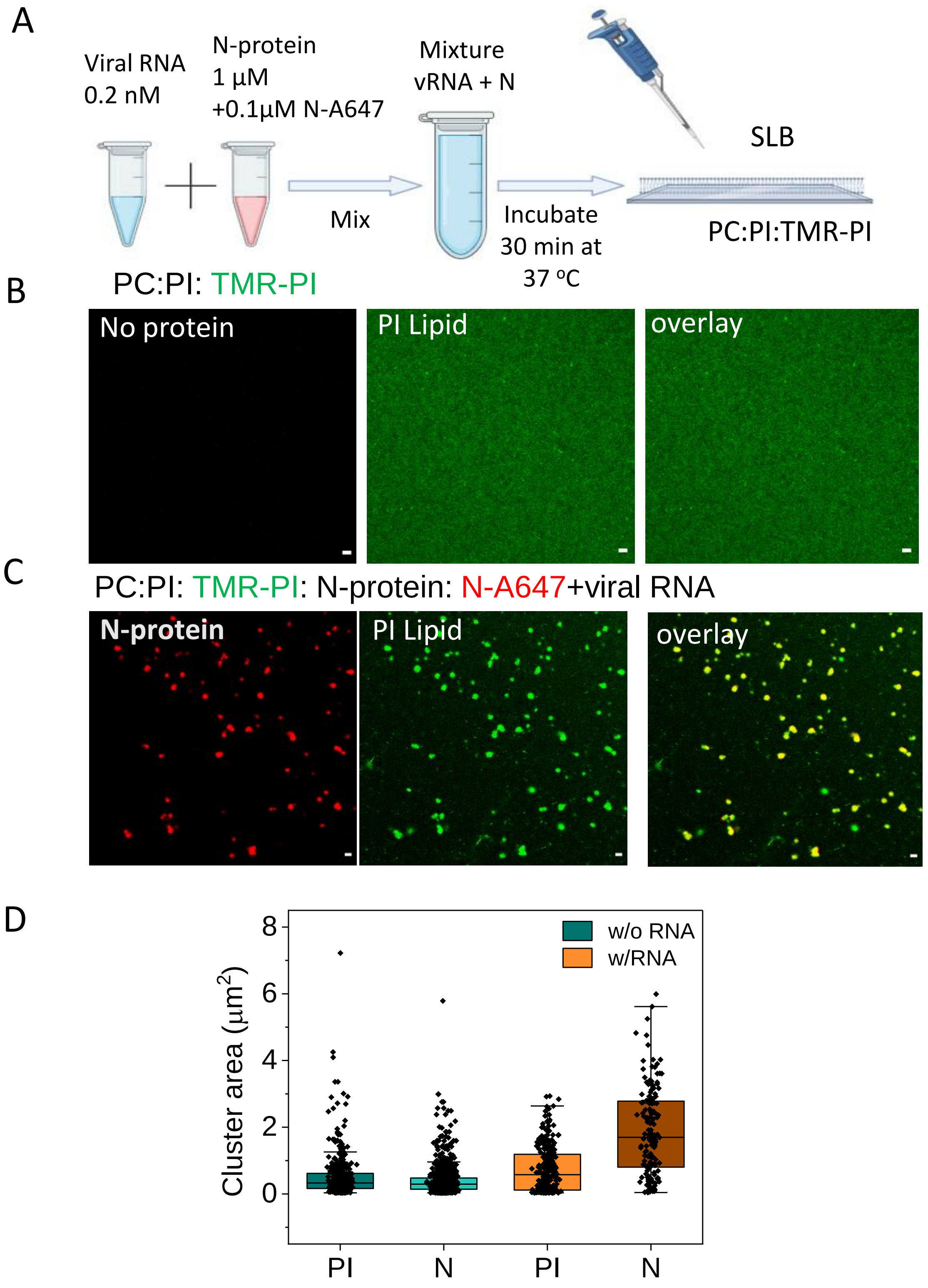


**Supplemental Figure 4: PI-TMR lipid clustering by N/RNP on simple PC:PI biomimetic membrane using fluorescence confocal microscopy. (A)** Scheme of experimental procedure.

(**B**) Confocal images of SLB made of [PC:PI 80:20] with 1% TMR-PI, N and PI clusters induced by N are visible on the SLBs. Scale bar is XX. Control is the image with No protein of the SLB containing the TMR-PI lipid and in buffer solution before adding the N protein. (**C**) The N protein was labelled with 1-5% N-A647 and mixed to unlabelled N prior to incubation on SLBs for 30min at 37°C. (**D**) Graph showing comparative measurements of PI and N cluster sizes with and without viral RNA. Data measurements are the Mean with a dot-box showing 25 to 75% of the clusters issued from 2 independent pooled experiments. Each black dot is one PI or N clusters (n= 200-600 clusters per condition).

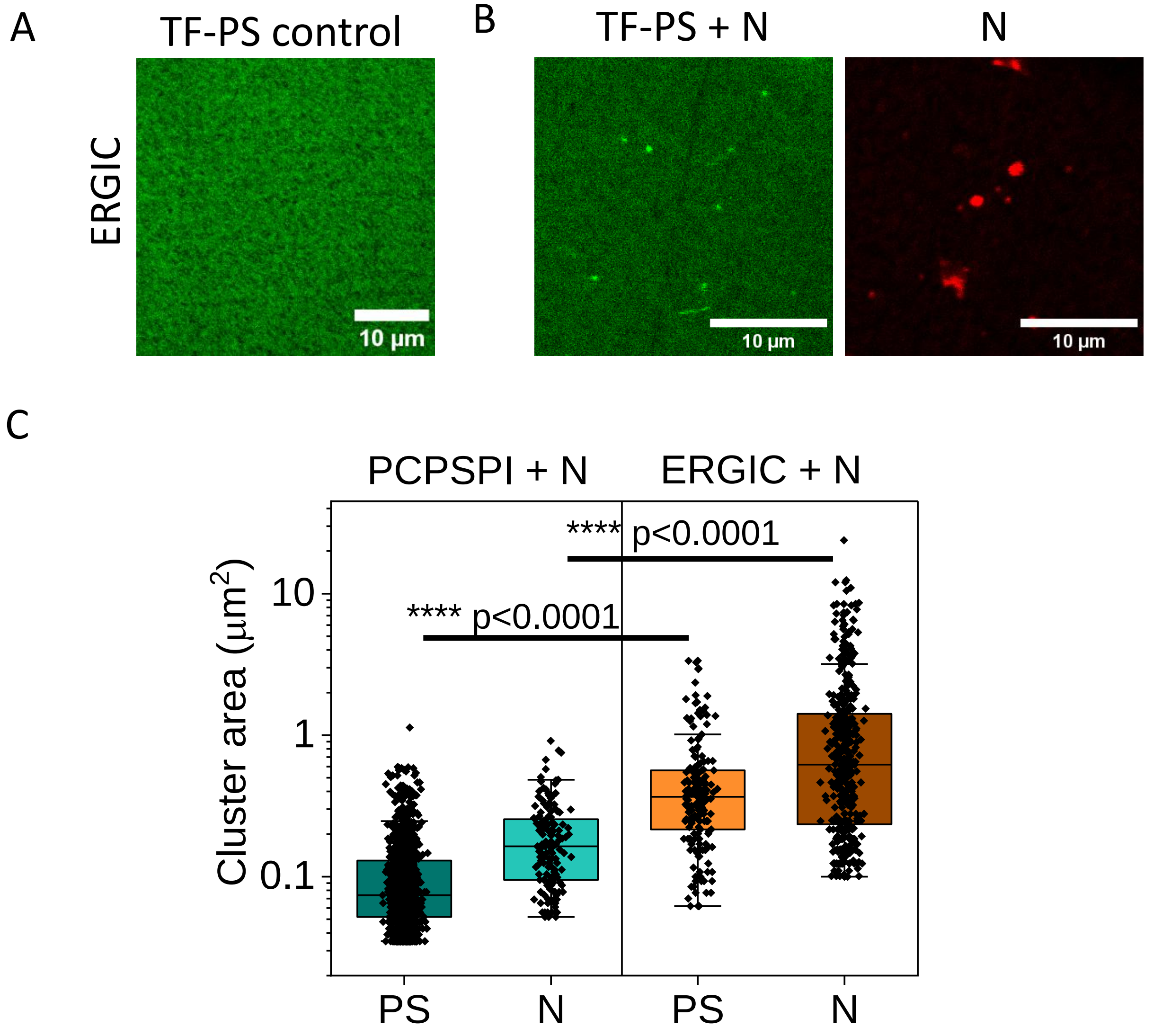


**Supplemental Figure 5: Comparing TF-PS lipid clustering by the N protein on simple and ERGIC biomimetic membranes. (A-B)** Confocal images of ERGIC-like composition made of [PC:PE:PS:PI:Chol (50:20:10:10:10)] with TopFluor-PS without N (A), and with N (B): N and PS clusters induced by N are visible on the SLBs. Scale bar is 10µm. PI lipid was labelled with TF-PI (0.5%). Control is the image of the SLB containing the TF-PS lipid in buffer solution before adding the N protein. The N protein was labelled with 1-5% N-A647 and mixed to unlabelled N prior to incubation on SLB for 30min at 37°C. **(C)** Graph showing comparative measurements of PS and N cluster sizes on simple PC:PS:PI (80:10:10) and ERGIC membranes. Data measurements are the Mean with a dot-box showing 25 to 75% of the clusters issued from 2 independent pooled experiments. Each black dot is one PS or N cluster (n= 140 to 1300 clusters per condition).

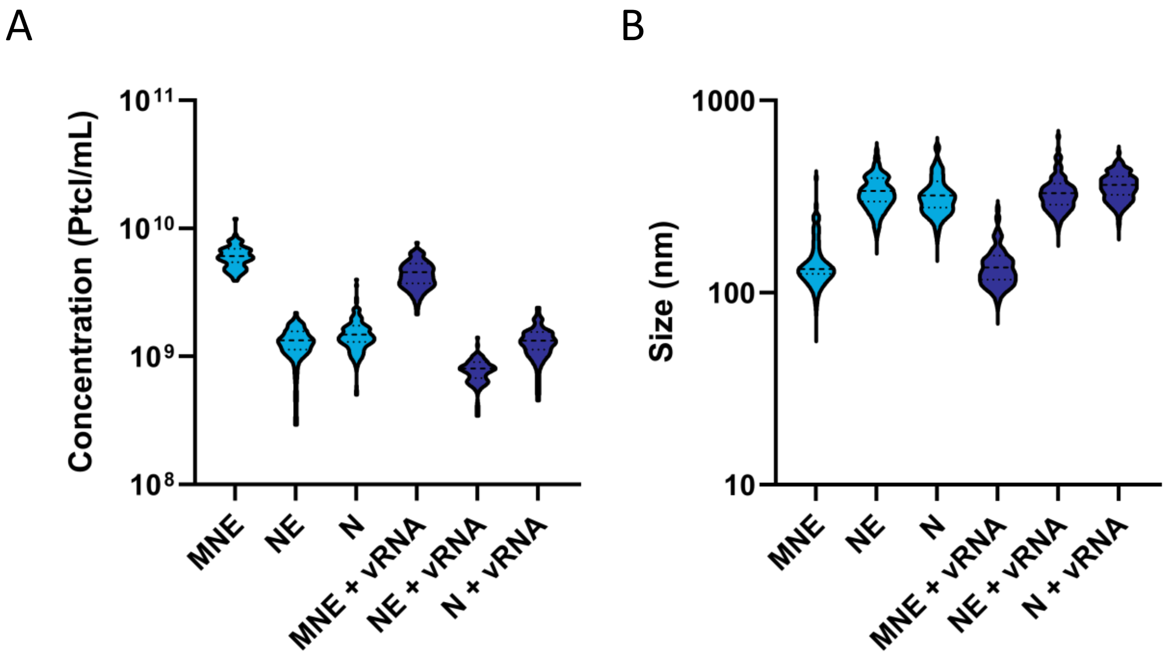


**Supplemental Figure 6: Concentration and size of MNE/N-mEOS2 Virus-like Particles using Nanoparticle Tracker Analyzer.** Purified particles MN/NmEOS2E or N/NmEOS2or N/NmEOS2 E produced by transfected 293THEK cells were injected into the ZetaView nanoparticle tracking analyzer. (A) The concentration of fluorescent particles was measured for each sample, giving a distribution; the peak of each distribution was then compiled into a violin plot. (B) Their hydrodynamic size was similarly measured and plotted in a second violin plot. ~70 fields of view corresponding to ~3000 particles per condition were measured.
